# Visual discrimination of optical material properties: a large-scale study

**DOI:** 10.1101/800870

**Authors:** Masataka Sawayama, Yoshinori Dobashi, Makoto Okabe, Kenchi Hosokawa, Takuya Koumura, Toni Saarela, Maria Olkkonen, Shin’ya Nishida

## Abstract

Complex visual processing involved in perceiving the object materials can be better elucidated by taking a variety of research approaches. Sharing stimulus and response data is an effective strategy to make the results of different studies directly comparable and can assist researchers with different backgrounds to jump into the field. Here, we constructed a database containing several sets of material images annotated with visual discrimination performance. We created the material images using physically-based computer graphics techniques and conducted psychophysical experiments with them in both laboratory and crowdsourcing settings. The observer’s task was to discriminate materials on one of six dimensions (gloss contrast, gloss distinctness-of-image, translucent vs. opaque, metal vs. plastic, metal vs. glass, and glossy vs. painted). The illumination consistency and object geometry were also varied. We used a non-verbal procedure (an oddity task) applicable for diverse use-cases such as cross-cultural, cross-species, clinical, or developmental studies. Results showed that the material discrimination depended on the illuminations and geometries and that the ability to discriminate the spatial consistency of specular highlights in glossiness perception showed larger individual differences than in other tasks. In addition, analysis of visual features showed that the parameters of higher-order color texture statistics can partially, but not completely, explain task performance. The results obtained through crowdsourcing were highly correlated with those obtained in the laboratory, suggesting that our database can be used even when the experimental conditions are not strictly controlled in the laboratory. Several projects using our dataset are underway.

## Introduction

Humans can visually recognize a variety of material properties of the objects they daily encounter. Although material properties, such as glossiness and wetness, substantially contribute to recognition, the contributions of value-based decision making, motor control, and computational and neural mechanisms underlying material perception had been overlooked until relatively recently—for a long time vision science mainly used simple artificial stimuli to elucidate the underlying brain mechanisms. In the last two decades, however, along with the advancement in computer graphics and machine vision, material perception becomes one of major topics in vision science (Adelson, 2001; Fleming, 2017; Nishida, 2019).

Visual material perception can be considered to be an estimation of material-related properties from an object image. For example, gloss/matte perception entails a visual computation of the diffuse and specular reflections of the surface. However, psychophysical studies have shown that human gloss perception does not have robust constancy against changes in surface geometry and illumination (e.g., Nishida & Shinya, 1998; Fleming et al. 2003), the other two main factors of image formation. Such estimation errors have provided useful information as to what kind of image cues humans use to estimate gloss. A significant number of psychophysical studies have been carried out not only on gloss, but also on other optical material properties (e.g., translucency, transparency and wetness) (Fleming et al., 2005; Motoyoshi, 2010; Xiao et al., 2014; Sawayama, Adelson, & Nishida, 2017, Liao et al., 2021) and mechanical material properties (e.g., viscosity, elasticity) (Kawabe et al., 2015; Paulun et al., 2017; van Assen, Barla & Fleming, 2018, van Assen, Nishida, & Fleming, 2020). Neurophysiological and neuroimaging studies have revealed various neural mechanisms underlying material perception (Kentridge et al., 2012; Nishio et al., 2012, 2014; Miyakawa et al., 2017). Some recent studies have also focused on developmental, environmental, and clinical factors of material processing (Yang et al., 2015; Goda et al., 2016; Ohishi et al. 2018). For instance, Goda et al. (2016) showed in their monkey fMRI study that the visuo-haptic experience of material objects alters the visual cortical representation. In addition, large individual differences in the perception of colors and materials depicted in one photo (#TheDress) has attracted a broad range of interest and has provoked intensive discussions (Brainard & Hurlbert, 2015; Gegenfurtner et al., 2015).

A promising strategy for a more global understanding of material perception is to promote multidisciplinary studies comparing behavioral/physiological responses of humans and animals obtained under a variety of developmental, environmental, cultural, and clinical conditions. There are two problems however. One lies in the high degree of freedom in selecting experimental stimulus parameters and task procedures. Since the appearance of a material depends not only on reflectance parameters, but also on geometry and illumination, all of which are high dimensional, use of different stimuli (and different tasks) in different studies could impose serious limitations on direct data comparisons. The other problem is the technical expertise necessary for rendering realistic images, which could discourage researchers unfamiliar with graphics from starting material perception studies.

Aiming at removing these obstacles, we attempted to build a database that can be shared among multidisciplinary material studies. We rendered several sets of material images. The images in each set were changed in one of material dimensions in addition to illumination and viewing conditions. We then measured the behavioural performance for those image sets using a large number of “standard” observers. We used a simple task that can be used in a variety of human, animal and computational studies. By using our database, one would be able to efficiently start a new study, shortening time for stimulus preparation, as well as time for control data collection with standard human observers.

Specifically, we selected six dimensions of material property (Fig. 1). These dimensions have been extensively studied in the past material perception studies. Most of them can be unambiguously manipulated by changing the corresponding rendering parameters. Although we attempted to cover a wide range of optical material topics, we never believe this an exclusive list of critical material properties vision science should challenge. Our intention is not to build the standard database for all material recognition research, but to make one primitive test set that promotes further examination of the previous findings on material recognition in more diverse research contexts. (see Discussion).

**Figure 1.**
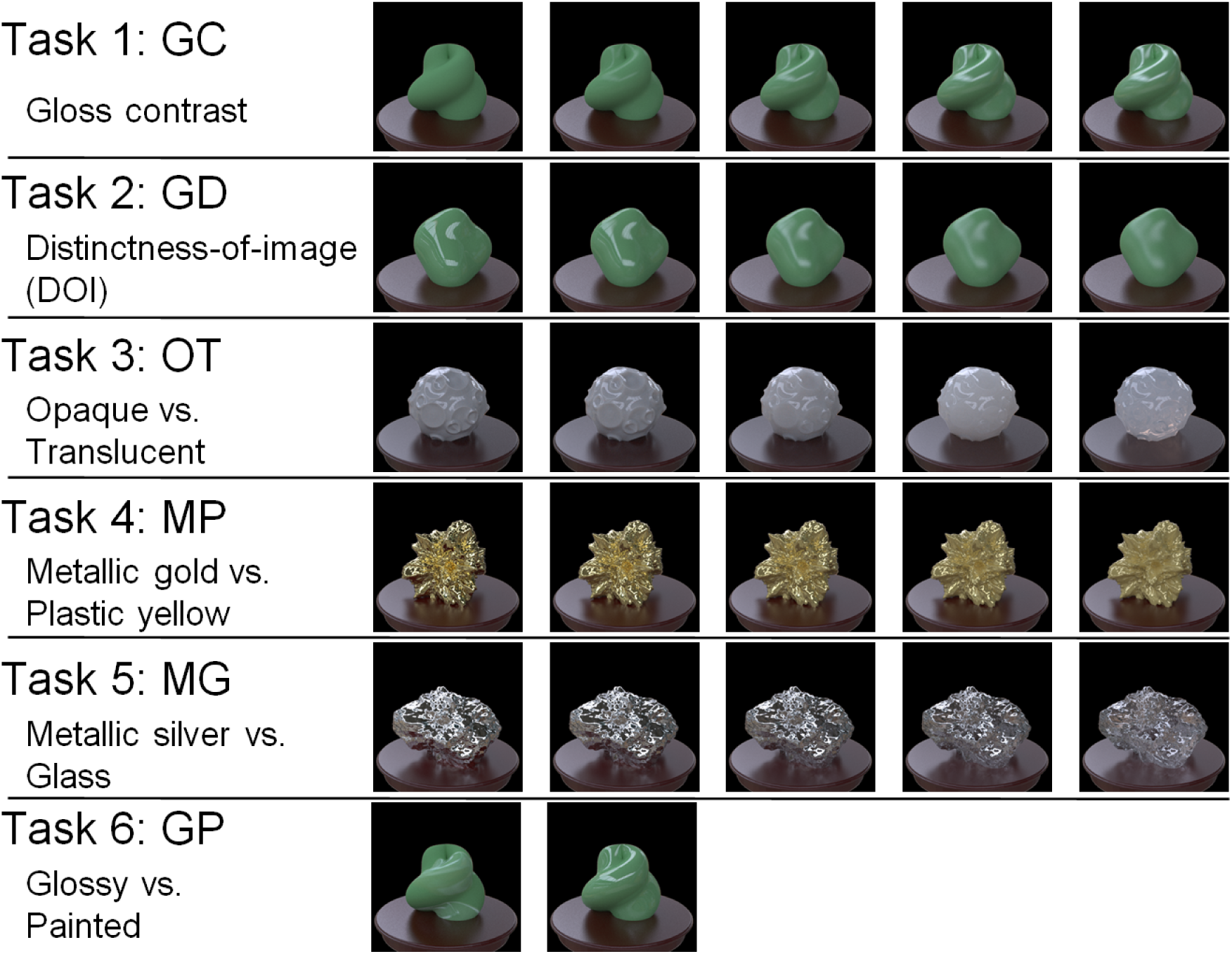
Schematic overview of six tasks recorded in the database.

Three of these dimensions are related to gloss (Fig. 1, Task 1: GC, Task 2: GD, and Task 6: GP), the most widely investigated material attribute (Pellacini et al., 2000; Fleming et al., 2003; Motoyoshi et al., 2007; Olkkonen & Brainard, 2010; Doerschner et al., 2011; Kim et al., 2011; Marlow et al., 2011; 2012; Kentridge et al., 2012; Sun et al., 2015; Nishio et al., 2014; Adams et al., 2016; Miyakawa et al., 2017; Storrs et al., 2021; Schmid et al., 2021). We controlled the contrast gloss and distinctness-of-image (DOI) gloss (gloss distinctness-of-image) as in previous studies (Pellacini et al., 2000; Fleming et al., 2003; Nishio et al., 2014). For instance, Nishio et al. (2014) found neurons in the inferior temporal cortex (ITC) of monkeys that selectively and parametrically respond to gloss changes in these two dimensions. We also controlled the spatial consistency of specular highlights, which is another stimulus manipulation of gloss perception (Fig. 1, Task 6: GP). By breaking the spatial consistency, some highlights look like albedo changes by white paint (Beck & Prazdny, 1981; Kim et al., 2011; Marlow et al., 2011; Sawayama & Nishida, 2018). Besides gloss perception, translucency perception has also been widely investigated (Fleming & Bülthoff, 2005; Motoyoshi, 2010; Nagai et al., 2013; Gkioulekas et al., 2013; Xiao et al., 2014; Chadwick et al., 2018; Gigilashvili et al., 2021). We adopted the task of discriminating opaque from translucent objects by controlling the thickness of the translucent media (Fig. 1, Task 3: OT). Furthermore, we adopted the task of plastic-yellow/gold discrimination (Okazawa et al., 2011, Task 4: MP) and glass/silver discrimination (Kim & Marlow, 2016; Tamura et al., 2019, Task 5: MG).

We used an oddity task (Fig. 3) to evaluate the capability of discriminating each material dimension. We chose this task because it requires neither complex verbal instruction, nor verbal responses by the observer. Therefore, it can be applied to a wide variety of observers including infants, animals, and machine vision algorithms, and their task performances can be directly compared. Indeed, several research projects using our dataset are underway (see the Discussion section).

To control the task difficulty, we varied the value of the parameter of each material dimension. In addition, we manipulated the stimulus in two ways that affected the task difficulty. First, we set three illumination conditions: one set of stimuli included images of different poses taken in identical illumination environments (Fig. 2a, Illumination condition 1); the second set contained stimuli of identical poses taken in slightly different illumination environments (Fig. 2a, Illumination condition 2); the third set contained identical poses taken in largely different illumination environments (Fig. 2a, Illumination condition 3). Second, we used the five different object geometries for each task (Fig. 2b).

**Figure 2.**
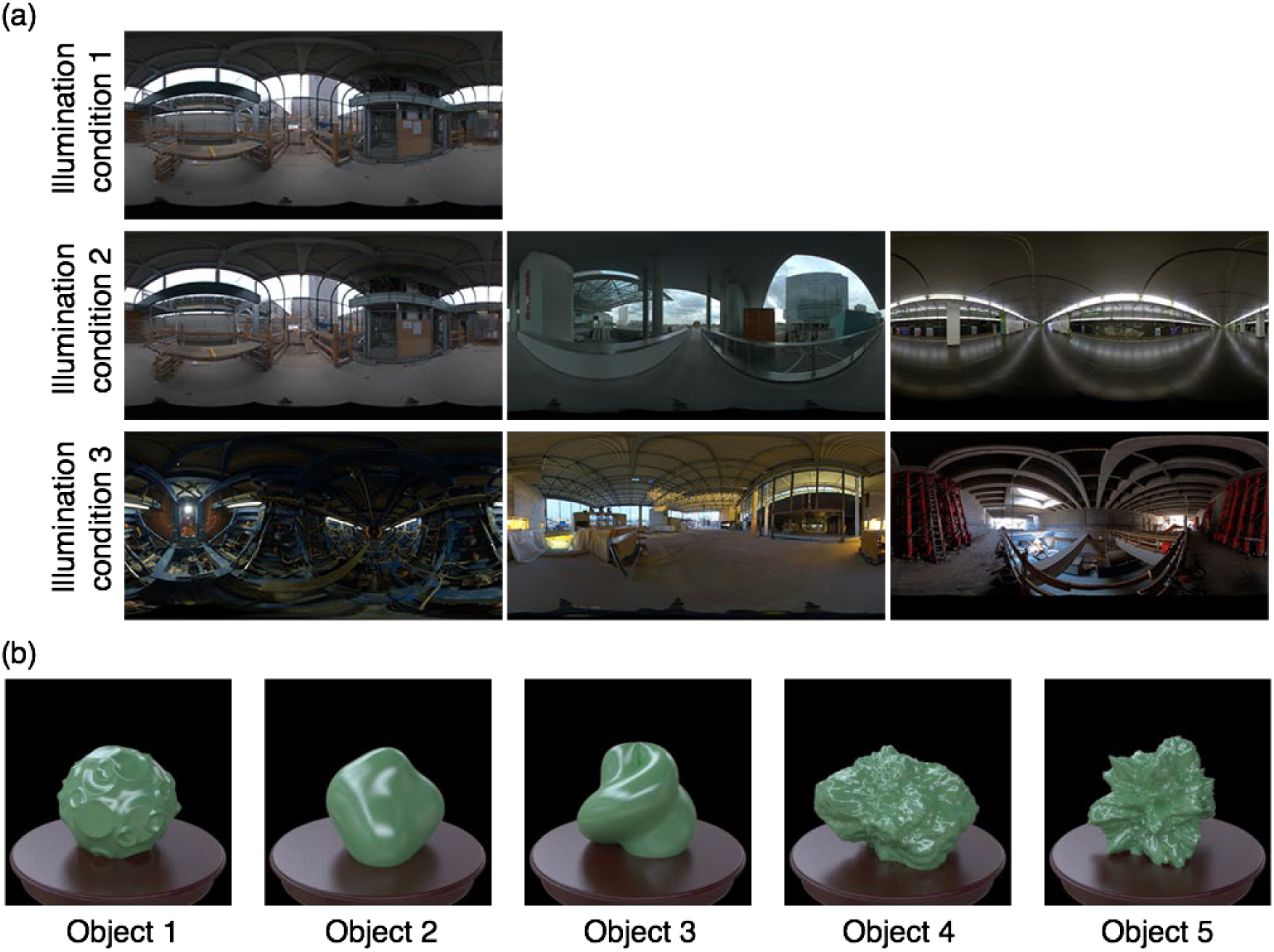
(a) Illumination conditions. Object images were rendered with six global illumination environments and were presented to observers under three illumination conditions. Under illumination condition 1, a stimulus display consisted of four objects (same shape, different poses) rendered with the same illumination environment. Under illumination condition 2, a stimulus display consisted of three objects (same shape, same pose) rendered with slightly different (in terms of their pixel histograms) light probes. Under illumination condition 3, a stimulus display consisted of three objects (same shape, same pose) rendered with largely different illumination environments. (b) Geometrical conditions. We used five different object shapes for each material task under each illumination condition. The stimulus condition is also summarized in Table 1.

**Figure 3.**
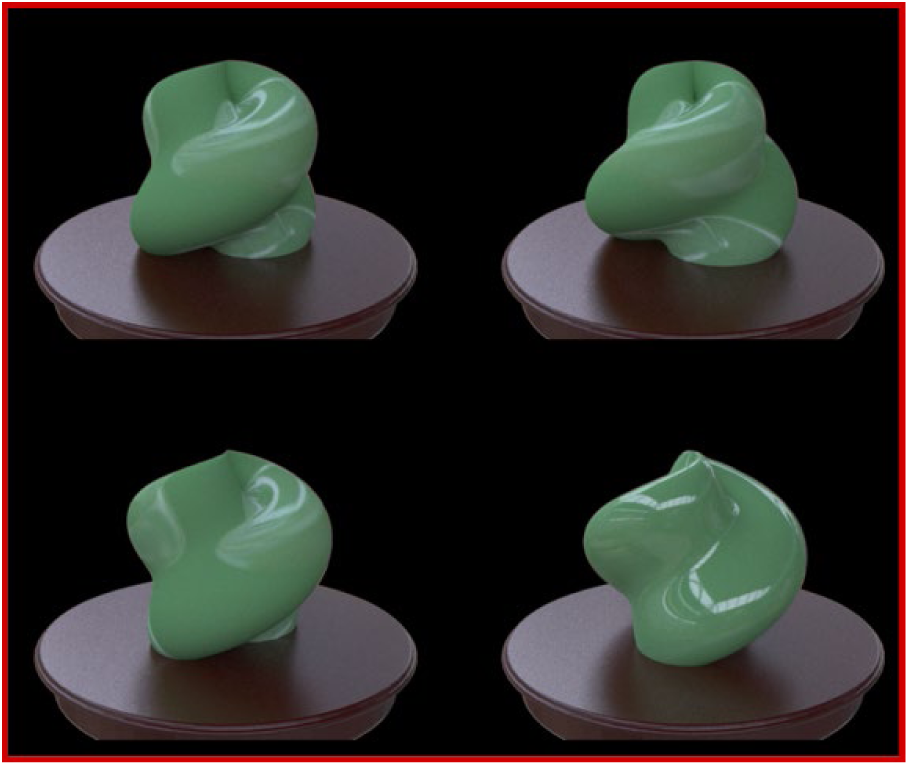
Example of a four-object oddity task (illumination condition 1) used for collecting standard observer data. The observers were asked to select which image was the odd one out in the four images. We did not tell the observer that the experiment was on material recognition. We conducted experiments both in the laboratory and through crowdsourcing.

**Table 1.**
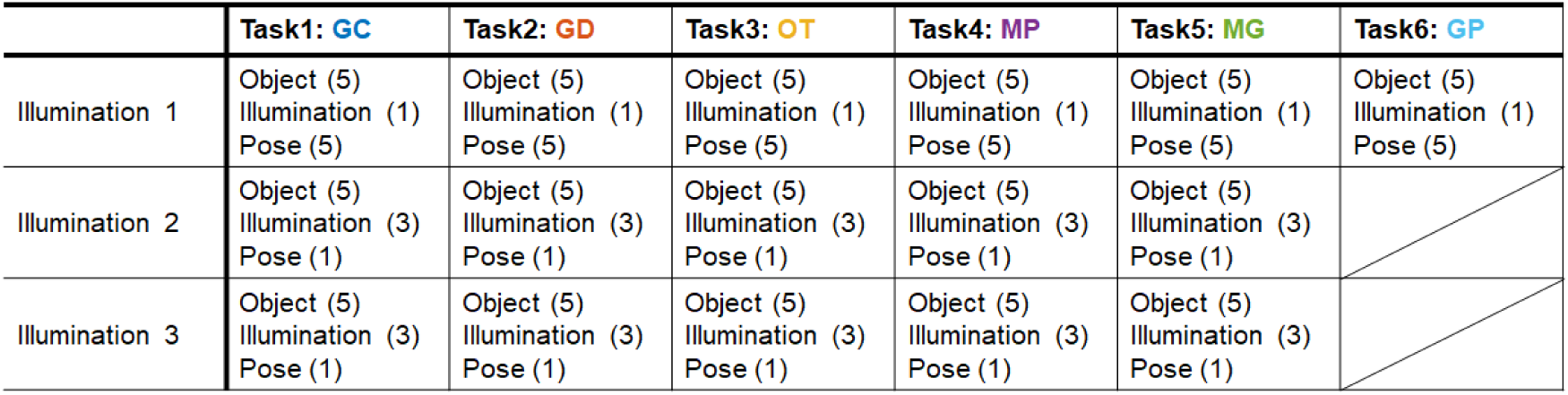
The summary of stimulus condition. The digit in parentheses indicates the number of each condition.

We wish to collect data from a large number of observers. A laboratory experiment affords control over the stimulus presentation environment, but is unsuited to collecting a large amount of data from numerous participants. In contrast, one can collect a lot of data through crowdsourcing, at the expense of reliable stimulus control. To overcome this trade-off, we conducted identical psychophysical experiments both in the laboratory and through crowdsourcing. This enabled us to evaluate individual difference distributions along with the effects of environmental factors on task performance.

In sum, we made a large set of image stimuli for evaluations of visual discrimination performance on six material dimensions (gloss contrast, DOI (distinctness-of-image) of gloss, translucency-opaque, plastic-gold, glass-silver and glossy-painted) and measured a large number of adult human observers performing oddity tasks in the laboratory and through crowdsourcing. The tasks had three illumination conditions and five object geometries. Although the original motivation of this project was to make a standard stimulus-response dataset of material recognition for promotion of multidisciplinary studies, it also has its own scientific value as it is the first systematic comparison of the effects of illumination condition and object geometry, as well as of individual variations across a variety of material dimensions. Our data include several novel findings, as shown below.

## Methods

We evaluated the observers’ performance of six material recognition tasks. We selected such tasks that had been used in previous material studies: 1) Contrast gloss discrimination (GC); 2) DOI (distinctness-of-image) discrimination (GD); 3) Opaque vs. translucent (OT); 4) Metallic gold vs. plastic yellow (MP); 5) Metallic silver vs. glass (MG); 6) Glossy vs. painted (GP). For each task, we used five geometry models. We used six global illuminations for tasks 1-5 and one for task 6. We conducted behavioral experiments using an oddity task, which can be used even with human babies, animals, and brain-injured participants, because it does not entail complex verbal instructions. In the experiment, the observers were asked to select the stimulus that represented an oddity among three or four object stimuli. They were not given any feedback about whether their responses were correct or not. We controlled the task difficulty by changing the illumination and material parameters. To test the generality of the resultant database, we conducted identical experiments in the laboratory and through crowdsourcing.

### Image generation for making standard image database

We utilized the physically-based rendering software called *Mitsuba* (Jakob 2010) to make images of objects consisting of different materials, and we controlled six different material dimensions.

#### Material for tasks 1) Gloss discrimination (contrast dimension) (Task 1: GC) and 2) Gloss discrimination (DOI dimension) (Task 2: GD)

To control the material property of the gloss discrimination tasks, we used the perceptual light reflection model proposed by Pellacini et al. (2000). They constructed a model based on the results of psychophysical experiments using stimuli rendered by the Ward reflection model (Ward, 1992) and rewrote the Ward model parameters in perceptual terms. The model of Pellacini et al. has two parameters, named *d* and *c*, and they roughly correspond to the DOI gloss and the contrast gloss of Hunter (1937). The difficulty of our two gloss discrimination tasks was controlled by separately modulating these two parameters.

The parameter space of the Ward reflection model can be described as follows.

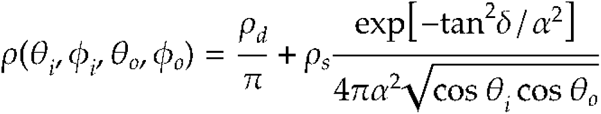

where ρ(θi,φi,θo,φo) is the surface reflection model, and θi, φi, and θo, φo are the incoming and outgoing directions, respectively. The model has three parameters; ρd is the diffuse reflectance of a surface, ρs is the energy of its specular component, and α is the spread of the specular lobe. Pellacini et al. (2000) defined two perceptual dimensions, *c* and *d* on the basis of the Ward model’s parameters. *d* corresponds to DOI gloss and is calculated from α, while *c* corresponds to perceptual glossiness contrast and is calculated from ρs and ρd, using the following formula:

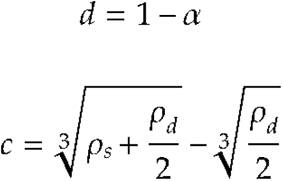

Although more physically feasible BRDF models than the Ward model have been proposed for gloss simulation (Ashikmin et al., 2000; Walter et al., 2007), we based ours on the Ward model because it has been used in many previous psychophysics and neuroscience studies (Nishio et al., 2014).

For the task of gloss discrimination in the contrast dimension, the specular reflectance ρs was varied in a range from 0.00 to 0.12 in 0.02 steps while keeping the diffuse reflectance ρd constant (0.416), indicating the contrast parameter: 0, 0.018, 0.035, 0.052, 0.067, 0.082, and 0.097. The distinctness-of-image d was the fixed value (0.94). (Fig. 4, Task 1: GC). As c gets closer to 0, the object appears to have a matte surface. The specular reflectance ρs of the non-target stimulus in the task was 0.06.

**Figure 4.**
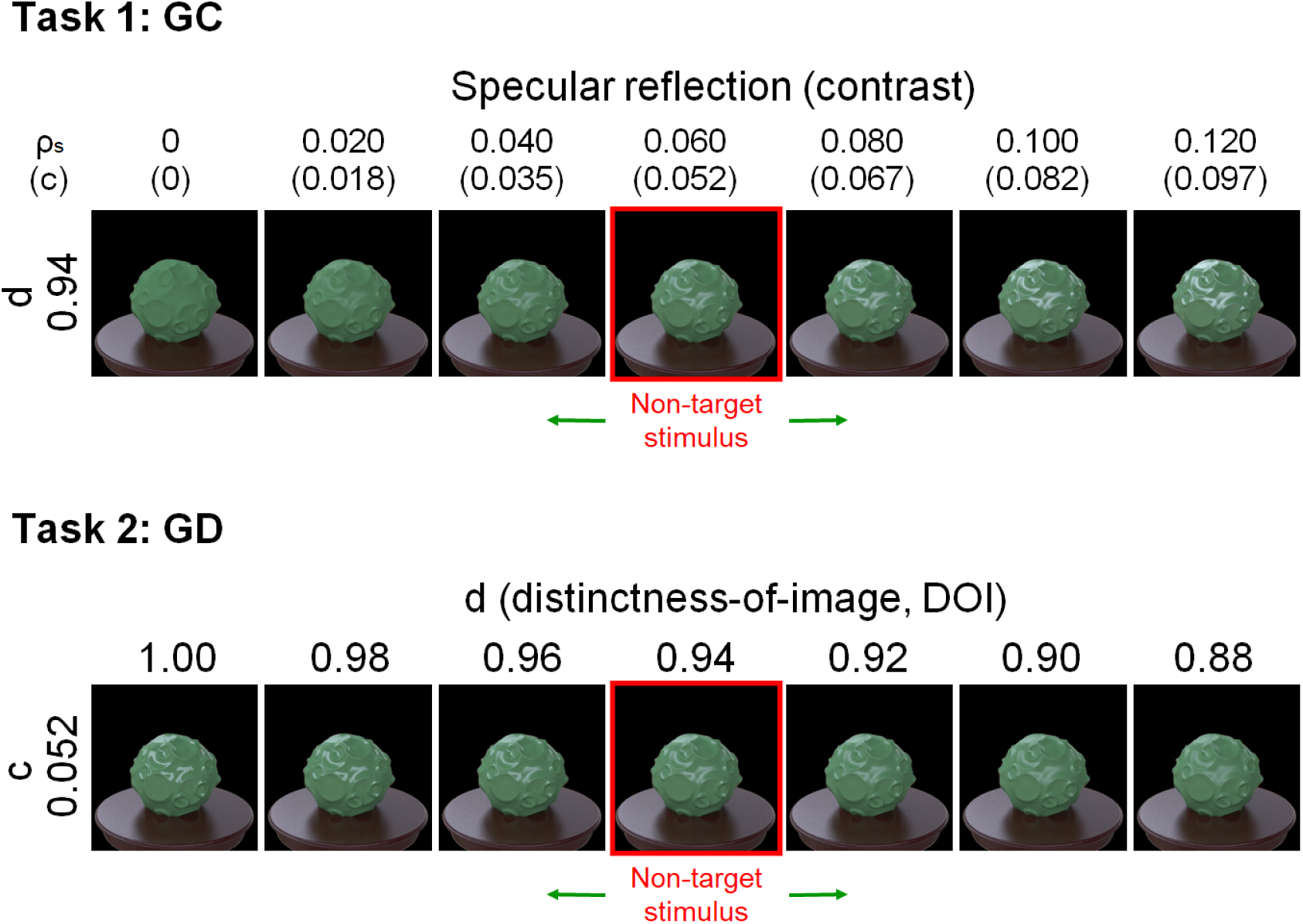
Material examples of tasks 1 (GC) and 2 (GD). For task 1 (GC), the specular reflectance of the odd target stimulus was varied from 0.00 to 0.12. The non-target stimuli that were presented as the context objects in each task had specular reflectance of 0.06. For task 2 (GD), the DOI parameter of the target specular reflection was varied from 1.00 to 0.88, while that of the non-target stimuli was 0.94.

For the experiment of gloss discrimination in the DOI dimension, the parameter *d* was varied from 0.88 to 1.00 in 0.02 steps while keeping ρs constant (0.06) (Fig. 4, *Task 2: GD*). As *d* gets closer to 1.00, the highlights of the object appear sharper. The DOI parameter, *d,* of the non-target stimuli was 0.94.

#### Material for task 3) Opaque vs. Translucent (Task 3: OT)

To make translucent materials, we used the function of homogeneous participating medium implemented in the Mitsuba renderer. In this function, a flexible homogeneous participating medium is embedded in each object model. The intensity of the light that travels in the medium is decreased by scattering and absorption and is increased by nearby scattering. The parameters of the absorption and scattering coefficients of the medium describe how the light is decreased. We used the parameters of the “Whole milk” measured by Jensen et al. (2001). The parameter of the phase function describes the directional scattering properties of the medium. We used an isotropic phase function. To control the task difficulty, we modulated the scale parameter of the scattering and absorption coefficients. The parameter describes the density of the medium. The smaller the scale parameter is, the more translucent the medium becomes. The scale parameter was varied as follows: 0.0039, 0.0156, 0.0625, 0.25, and 1.00 (Fig. 5, Task 3: OT). The scale parameter of the non-target stimulus in the task was 1.00. In addition, the surface of the object was modeled as a smooth dielectric material to produce strong specular highlights, as in previous studies (Gkioulekas, et al, 2013; Xiao et al., 2014). That is, non-target objects were always opaque, and the degree of transparency of the target object was changed.

**Figure 5.**
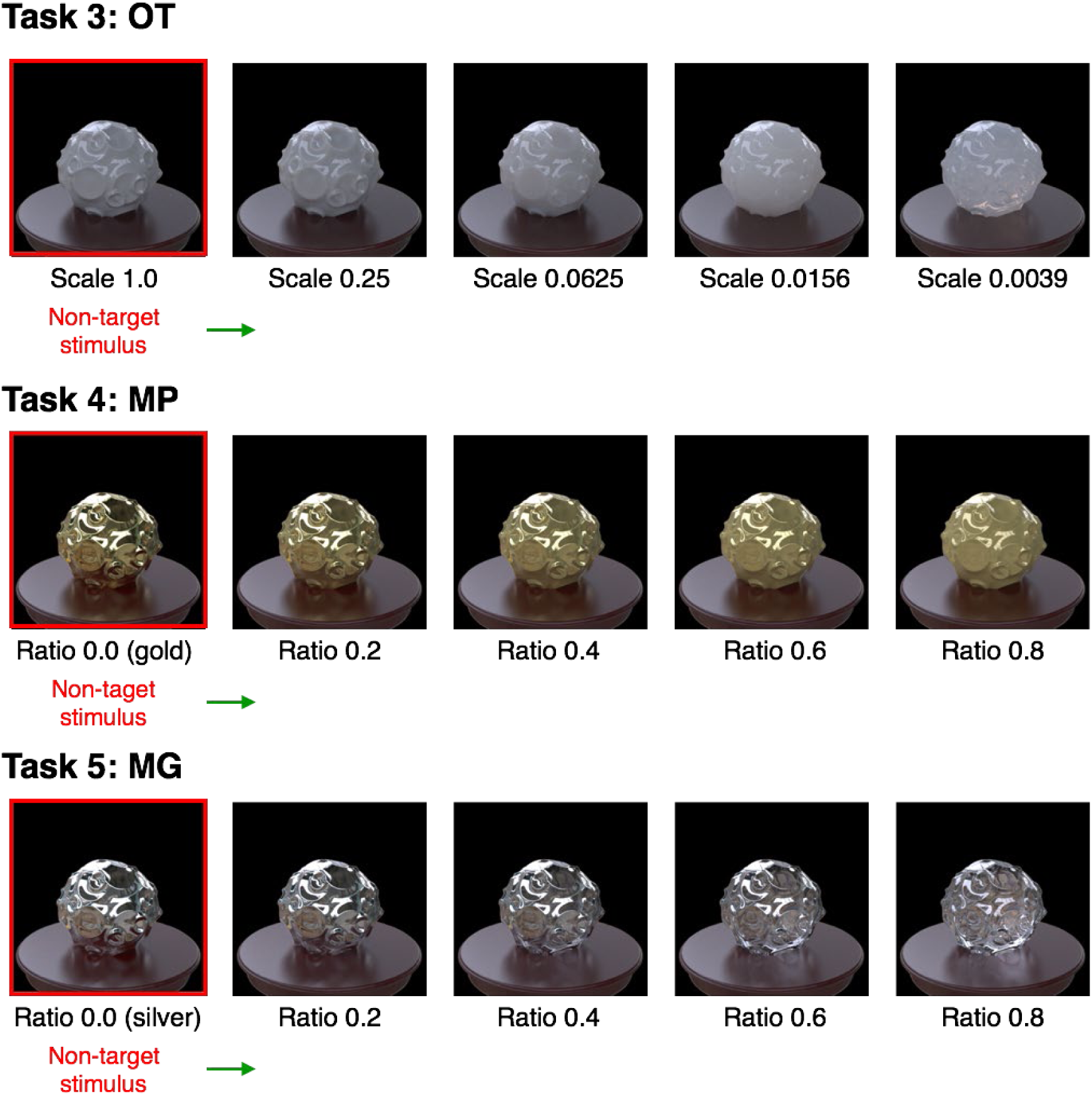
Material examples of tasks 3 (OT), 4 (MP), and 5 (MG). For task 3 (OT), the scale of the volume media that consisted of milk was varied from 1.0 to 0.0039. For task 4 (MP) and 5 (MG), the blending ratio of the two materials was varied from 0.0 to 0.8. The non-target stimuli in the tasks were shown as in the legend.

#### Material for task 4) Metallic gold vs. Plastic yellow (Task 4: MP)

To morph the material between gold and plastic yellow, we utilized a linear combination of gold and plastic BRDFs, which is implemented in the Mitsuba renderer. By changing the weight of the combination, the appearance of a material (e.g., gold) can be modulated toward that of the other material (e.g., plastic yellow). In this task, the weight was varied in a range from 0.00 to 0.80 in 0.20 steps (Fig. 5, Task 4: MP). The parameter of the non-target stimulus was 0, at which the material appeared to be pure gold.

#### Material for task 5) Metallic silver vs. Glass (Task 5: MG)

Similar to task 4), we utilized a linear combination of dielectric glass and silver materials, which is also implemented in the Mitsuba renderer. The weight of the combination was varied from 0.00 to 0.80. The parameter of the non-target stimulus was 0, at which the material appeared to be pure silver (Fig. 5, Task 5: MG).

As noted above, for Tasks 3, 4, and 5 in which the parameters of the target stimulus were varied between two material states (i.e., opaque vs. transparent, metallic vs. plastic, and metallic vs. glass), we placed the non-target objects at one end (i.e., one of two material states). If we placed the non-target stimuli in the middle of the stimulus variable as in Tasks 1 and 2, and when the difference between the target and non-target stimuli was small, the display only contained ambiguous material objects. In such cases, the observers might not pay attention to the material dimension relevant to the task. By placing the non-target at one extreme value, we could make the stimulus display always contain the object images in a specific material state, helping participants focus on the task relevant material dimension.

#### Material for task 6) Glossy vs. Painted (Task 6: GP)

The skewed intensity distribution due to specular highlights of an object image can be a diagnostic cue for gloss perception (Motoyoshi et al., 2007). However, when the specular highlights are inconsistent in terms of their position and/or orientation with respect to the diffuse shading component, they look more like white blobs produced by surface reflectance changes even if the intensity distribution is kept constant (Beck & Prazdny; 1981; Anderson & Kim, 2009; Kim et al., 2011; Marlow et al., 2011; Sawayama & Nishida, 2018). For our last task of glossy objects vs. matte objects with white paint, we rendered the glossy objects on the basis of Pellacini et al. (2000)’s model. The parameter *c* was set to 0.067, and the parameter *d* ranged from 0.88 to 1.00 in 0.04 steps (Fig. 6, lower). Considering material naturalness, these objects may not be typically encountered in the real world, but this task is theoretically important because it will provide insights into the underlying visual computation of material recognition.

**Figure 6.**
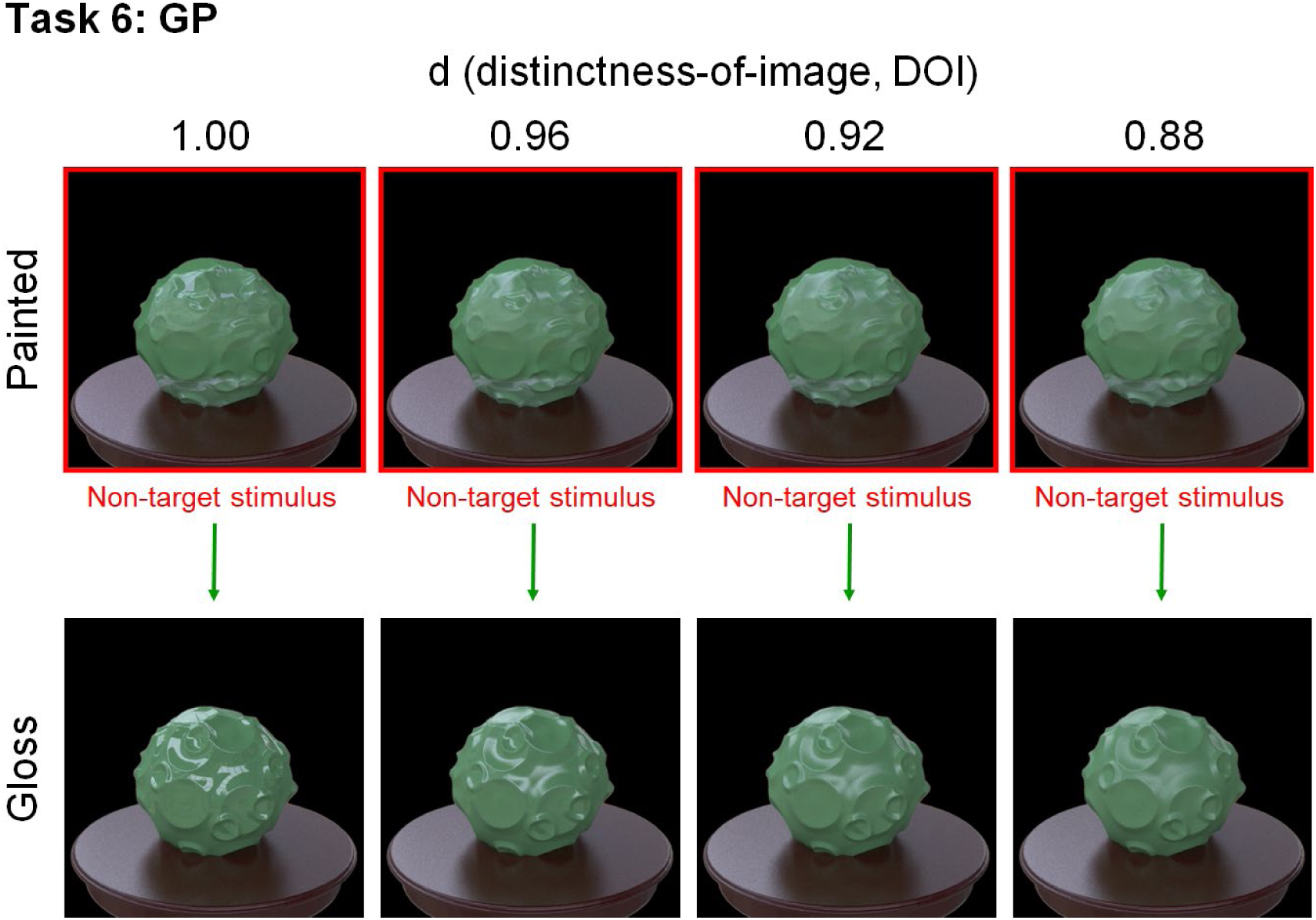
Material examples of task 6. The distinctness-of-image of the specular reflection was varied from 1.00 to 0.88. This parameter was the same for the non-target painted objects and the target glossy object in each stimulus display.

To make object images with inconsistent highlights (white paints), we rendered each scene twice with different object materials with identical shapes. First, we rendered a glossy object image by setting the diffuse reflectance to 0, i.e., the image that includes only specular highlights. The rendered image of specular highlights was a 2D texture for the second rendering. We eliminated the brown table when rendering the first scene. Next, we rendered a diffuse object image, i.e., one without specular reflection, with the texture of specular highlights. The object and illumination for the first and second renderings were the same. We mapped the specular image rendered in one object pose to the 3D geometry by a spherical mapping with repeating the image. Since the position of texture mapping was randomly determined, the highlight texture positions were inconsistent with diffuse shadings. We varied the parameter *d* of the first rendering from 1.00 to 0.88 (Fig. 6, lower). After we rendered the inconsistent-highlights image, the color histogram of the image was set to that of a consistent glossy object image by using a standard histogram matching method (Sawayama & Nishida, 2018).

We made task 6 only under Illumination 1. This is because it was hard to match the color distributions of the target and non-target stimuli for Illuminations 2 and 3, where one stimulus set was rendered under different illuminations. If we match the objects’ color histograms under these conditions, the object’s colors could be incongruent with their background colors (i.e., the table and the shadow in this scene). This could produce another cue to find an outlier, which making these conditions inappropriate for the task purpose.

#### Geometry

For each material, we rendered the object images by using five different abstract geometries (Fig. 2b). These geometries were made from a sphere by modulating each surface normal direction with different kinds of noise (see also ShapeToolbox: https://github.com/saarela/ShapeToolbox) (Saarela & Olkkonen, 2016, Saarela, 2018). Specifically, Object_1 was made from modulations of low-spatial-frequency noise and crater-like patterns. The source code of this geometry is available on the web (http://saarela.github.io/ShapeToolbox/gallery-moon.html). Object_2 was a bumpy sphere modulated by low-pass band-pass noise. Object_3 was a bumpy sphere modulated by sine-wave noise. Object_4 and Object_5 were bumpy spheres modulated by Perlin noise. These objects were also rendered using Shapetoolbox.

Five samples were too small to systematically vary shape parameters. Instead, we handcrafted sphere-based abstract shapes in such a way expected to maximize the shape diversity. It is known that even when rendering with the same reflectance function (BRDF), objects with smooth/low-frequency surface modulations and those with spiky/high-frequency surface modulations could have very different material appearance (Shinya & Nishida, 1998, Vangorp, Laurijssen, & Dutré, 2007). We therefore created five geometries with a variety of low and high spatial frequency surface modulations to see human material perception under widely different geometry conditions.

#### Illumination and pose

We used six high-dynamic-range (HDR) light-probe images as illuminations for rendering. These images were obtained from Bernhard Vogl’s light probe database (http://dativ.at/lightprobes/). To vary the task difficulty, we used three illumination conditions (illumination conditions 1, 2, and 3, Fig. 2a). Under illumination condition 1, the observers selected one oddity from four images in a task. We rendered the images by using an identical light probe (i.e., ‘Overcast Day/Building Site (Metro Vienna)’). We prepared five poses for each task of illumination condition 1 by rotating each object in 36-degree steps; four of them were randomly selected in each task.

Under illumination condition 2, the observers selected one oddity from three images in a task. We created the images by using slightly different (in terms of their pixel histograms) light probes (i.e., ‘Overcast Day/Building Site (Metro Vienna)’, ‘Overcast day at Techgate Donaucity’, and ‘Metro Station (Vienna Metro)’). The task procedure of illumination condition 3 was the same as that of illumination condition 2. For illumination condition 3, we created the three images by using light probes that were rather different from each other (‘Inside Tunnel Machine’, ‘Tungsten Light in the Evening (Metro Building Site Vienna)’, and ‘Building Site Interior (Metro Vienna)’). We computed the pixel histogram similarity for each illumination pair and used it as the distance for the multidimensional scaling analysis (MDS). We extracted three largely different light probes in the MDS space and used them for illumination condition 3. We also selected three similar light probes in the space and used them for illumination condition 2. The pose of each object in the illumination condition 2 and 3 was not changed. The stimulus condition is summarized in Table 1.

#### Rendering

To render the images, we used the integrator of the photon mapping method for tasks 1, 2, 4, 5, and 6 and used the integrator of the simple volumetric path tracer implemented in the Mitsuba renderer for task 3 (OT). The calculation was conducted using single-float precision. Each rendered image was converted into sRGB format with a gamma of 2.2 and saved as an 8-bit .png image.

### Behavioral experiments

#### Laboratory experiment

Twenty paid volunteers participated in the laboratory experiment. Before starting the experiment, we confirmed that all had normal color vision by having them take the Famsworth– Munsell 100 Hue Test and that all had normal or corrected-to-normal vision by having them take a simple visual acuity test. The participants were naïve to the purpose and methods of the experiment. The experiment was approved by the Ethical Committees at NTT Communication Science Laboratories.

The generated stimuli were presented on a calibrated 30-inch EIZO color monitor (ColorEdge CG303W) controlled with an NVIDIA video card (Quadro 600). Each participant viewed the stimuli in a dark room at a viewing distance of 86 cm, where a single pixel subtended 1 arcmin. Each object image of 512 x 512 pix was presented at a size of 8.5 x 8.5 degrees.

In each trial, four (Illumination 1) or three (Illumination 2 & 3) object images chosen for each task were presented on the monitor (Fig. 3). Measurements of different illumination conditions were conducted in different blocks. Under illumination condition 1, four different object images in different orientations were presented. Under illumination conditions 2 and 3, the three different object images had different illuminations. The order of illumination conditions 1, 2, and 3 was counterbalanced across observers. The observers were asked to report which of the object images looked odd by pushing one of the keys. The stimuli were presented until the observer made a response. The task instructions were simply to find the odd one with no further explanation about how it was different from the rest. The observers were not given any feedback about whether their response was correct or not. All made ten judgments for each task of illumination condition 1. Seventeen observers made ten judgments for each task of illumination condition 2, while three made only seven judgments due to the experiment’s time limitation. Seventeen observers made ten judgments for each task of illumination condition 3, while three made seven judgments due to the experiment’s time limitation.

#### Crowdsourcing experiment

In the web experiment, 416, 411, and 405 paid volunteers participated in the tasks of illumination conditions 1, 2, and 3, respectively. We recruited these observers through a Japanese commercial crowdsourcing service. All who participated under illumination condition 3 also participated under illumination conditions 1 and 2. Moreover, all who participated in illumination condition 2 had also participated under illumination condition 1. The experiment was approved by the Ethical Committees at NTT Communication Science Laboratories.

Each observer used his/her own PC’s or tablet’s web browser to participate in the experiment. We asked them to watch the screen from a distance of about 60 cm. Each object image was shown on the screen at a size of 512 x 512 pix. We didn’t strictly control the visual angle of the image participants observed.

The procedure was similar to that of the laboratory experiment. In each trial, four or three object images that had been chosen depending on the task were presented on the screen, as in Fig. 3. The measurement was conducted under illumination condition 1 first, followed by one under illumination condition 2 and one under illumination condition 3. The observers were asked to report which of the object images looked odd by clicking one of the images. Each participant made one judgment for each condition. The other steps of the procedure were the same as those in the laboratory experiment.

### Data analysis

For each oddity task, we computed the proportion that each participant got correct. The chance level of the correct proportion was 0.25 for illumination condition 1 and 0.33 for illumination conditions 2 and 3. We computed the sensitivity *d’* from each correct proportion by using a numerical simulation to estimate the sensitivity of the oddity task (Craven, 1992). We used the "Palamedes" data analysis library for the simulation (Kindom & Prins, 2010; 2016; Prins & Kingdom, 2018). To avoid values of infinity, we converted the one probability according to the total trial number (i.e., corrected the one value to 1-(1/2N), where N is the total trial number) in the simulation (Macmillan & Kaplan, 1985). For the laboratory experiment, we computed the sensitivity *d’* of each observer and averaged it across observers. For the crowdsourcing experiment, since each observer engaged in each task one time, we computed the proportion correct for each task from all observers’ responses and used it to compute *d’*.

## Results

In this section, we describe the results of our benchmark data acquisition. First, we evaluate the environment dependency of our experiment, the performance difference between the online and laboratory experiments. Then, we describe the illumination and geometry effect on each task. After discussing each task, we show how intermediate visual features contribute to task performance. In the end, we analyze the individual difference in each task.

### Environment dependence

For cross-cultural, cross-species, brain-dysfunction, and developmental studies, stimulus presentation on a monitor cannot always be strictly controlled because of apparatus or ethical limitations. Therefore, a performance validation of each task across different apparatuses is critical to decide which tasks the users of our database should select in their experimental environment. Figure 7a shows the results of the correlation analysis between the laboratory and crowdsourcing experiments. The coefficient of determination (R^2^) of the linear regression between the sensitivity d’ in the laboratory experiment and that of the crowdsourcing experiment is 0.79, indicating a high linear correlation. However, the slope of the regression is less than 1. This shows that the sensitivity of the crowdsourcing experiment was worse than that of the laboratory experiment, with many repetitions in general. These findings suggest that the present tasks maintain relative performance across different experimental environments.

**Figure 7.**
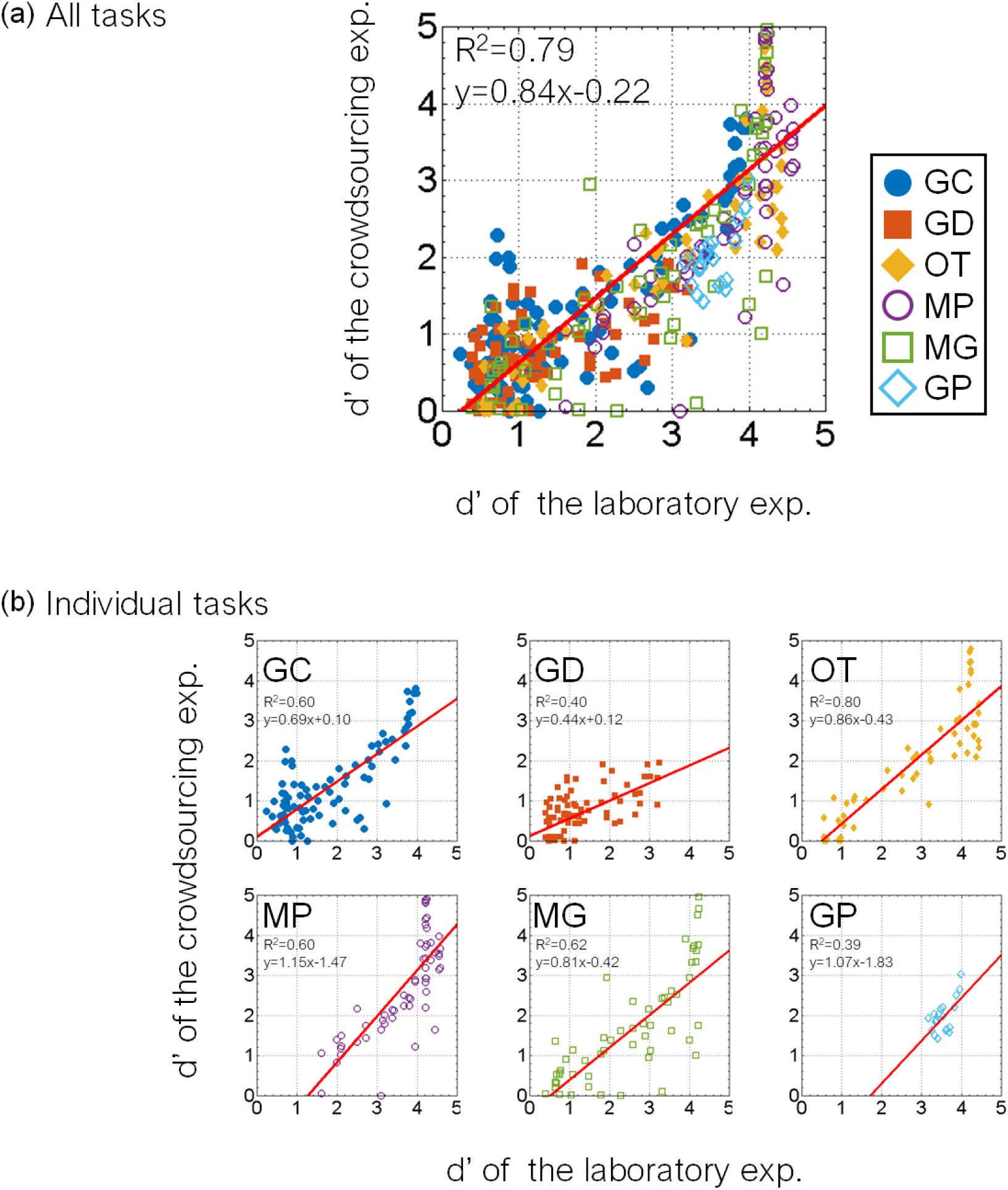
Results of laboratory and crowdsourcing experiments. The sensitivity d’ in each task in the crowdsourcing experiment is plotted as a function of that in the laboratory experiment. (a) Results of all tasks. Each plot indicates a task with an object, an illumination, and a difficulty. The red line indicates the linear regression between the crowdsourcing and laboratory results. The coefficient of determination (R^2^) of the regression and the equation are shown in the legend. The results show that the present tasks are generally robust across experimental environments. (b) Results of individual tasks. Different panels indicate tasks involving different materials. Each plot in a panel indicates a task with an object, illumination, and difficulty. The red line indicates the linear regression between the laboratory and crowdsourcing results. The coefficient of determination (R^2^) of the regression and the equation are shown in the legend. The accuracy of task 2 (GD) in the crowdsourcing experiment was generally lower than that in the laboratory experiment. The correlation of task 6 (GP) between the laboratory and crowdsourcing experiments was the worst.

Figure 7b shows the results for each task of the laboratory and crowdsourcing experiments in more detail. The coefficients of determination (R^2^) in tasks 1 to 6 are 0.60, 0.40, 0.86, 0.60, 0.62, and 0.39, respectively. The coefficient of task 6 (GP) was the worst, followed by task 2 (GD). As in the latter section, task 6 (GP) also showed large individual differences, and thus, the correlation between the laboratory and crowdsourcing experiments was decreased. The slope of the linear regression on task 2 (GD) was 0.44, and the proportion correct in the crowdsourcing experiment for tasks 2 were generally lower than those in the laboratory for tasks 2. In the laboratory experiment, we used a 30-inch LCD monitor, and the stimulus size of each image was presented at a size of 8.5 x 8.5 degrees, which we expected to be larger than when participants on the web observed the image on a tablet or PC. Task 2 (GD) is related to the distinctness-of-image of the specular reflection, and thus, the spatial resolution might have affected the accuracy of the observers’ responses, although the relative difficulty for task 2 (GD) even in the crowdsourcing experiment was similar to that in the laboratory experiment. These findings suggest that the absolute accuracy of task 2 (GD) depends largely upon the experimental environment.

### Illumination and geometry

Figures 8 to 13 show the performance of each task in the laboratory experiment. Different panels depict results obtained for different objects. Different symbols in each panel depict different illumination conditions. The results of the crowdsourcing experiment are shown in Appendix A. For task 1 to task 5 (Figures 8 to 12), we parametrically changed the material parameters, e.g., the contrast dimensions for task 1 (GC). Results show that the discrimination accuracy increased as the target material parameters deviated from the non-target one. This trend can be most evidently observed for Illumination 1 on each task condition. In contrast, the accuracy didn’t change much with the material parameters for some conditions. This trend can be observed on Illuminations 2 and 3 of task 1 (GC) and Objects 4 and 5 of task 2 (GD). For task 6, the relation of target and non-target stimuli is different from the other tasks. In this task, the non-target stimulus was made for each material parameter, i.e. the distinctness-of-image (DOI). As shown in Figure 13, this material parameter didn’t affect the task difficulty.

**Figure 8.**
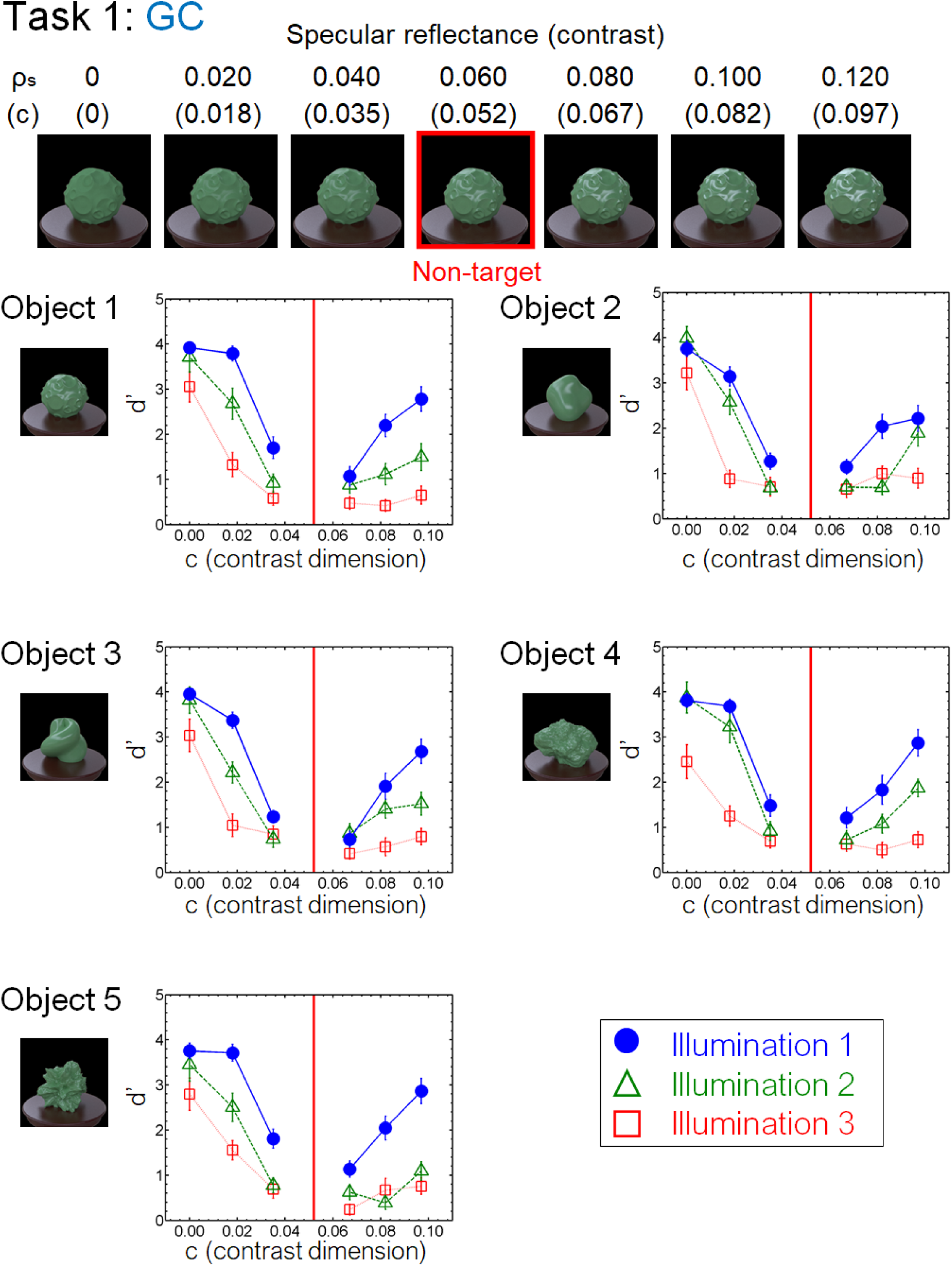
Results of task 1 (GC) in the laboratory experiment. Different panels show different objects. Different symbols in each panel depict different illumination conditions. The vertical red line in each panel indicates the parameter of the non-target stimulus. Error bars indicate ± 1 SEM across observers.

By comprehensively assessing material recognition performance across different stimulus conditions, we found novel properties that have been overlooked in the previous literature. One regards the geometrical dependence of material recognition. When object images changed in the gloss – distinctness-of-image dimension (task 2: GD, Fig. 9), the observers could detect the material difference better for smooth objects (Object 2 & 3) than for rugged objects (Object 4 & 5). In contrast, when the object images changed in the glossiness-contrast dimension (task 1: GC, Fig. 8), little geometrical dependence was found. We also found little geometrical dependence when observers detected highlight-shading consistency (task 6: GP, Fig. 13). While geometrical dependencies of glossiness perception have been reported before (Nishida & Shinya, 1998; Vangorp, Laurijssen, & Dutré, 2007), they were mainly about the effects of shape on apparent gloss characteristics, not on gloss discrimination. Furthermore, our results also show a geometrical dependence of translucency perception (task 3: OT, Fig. 10). Similar to the dependence on the distinctness-of-image dimension, the sensitivity changed between the smooth objects (Object 2 & 3) and rugged objects (Object 4 & 5), but in the opposite way. Specifically, the translucent difference was more easily detected for the rugged objects than for the smooth objects (Fig. 10).

**Figure 9.**
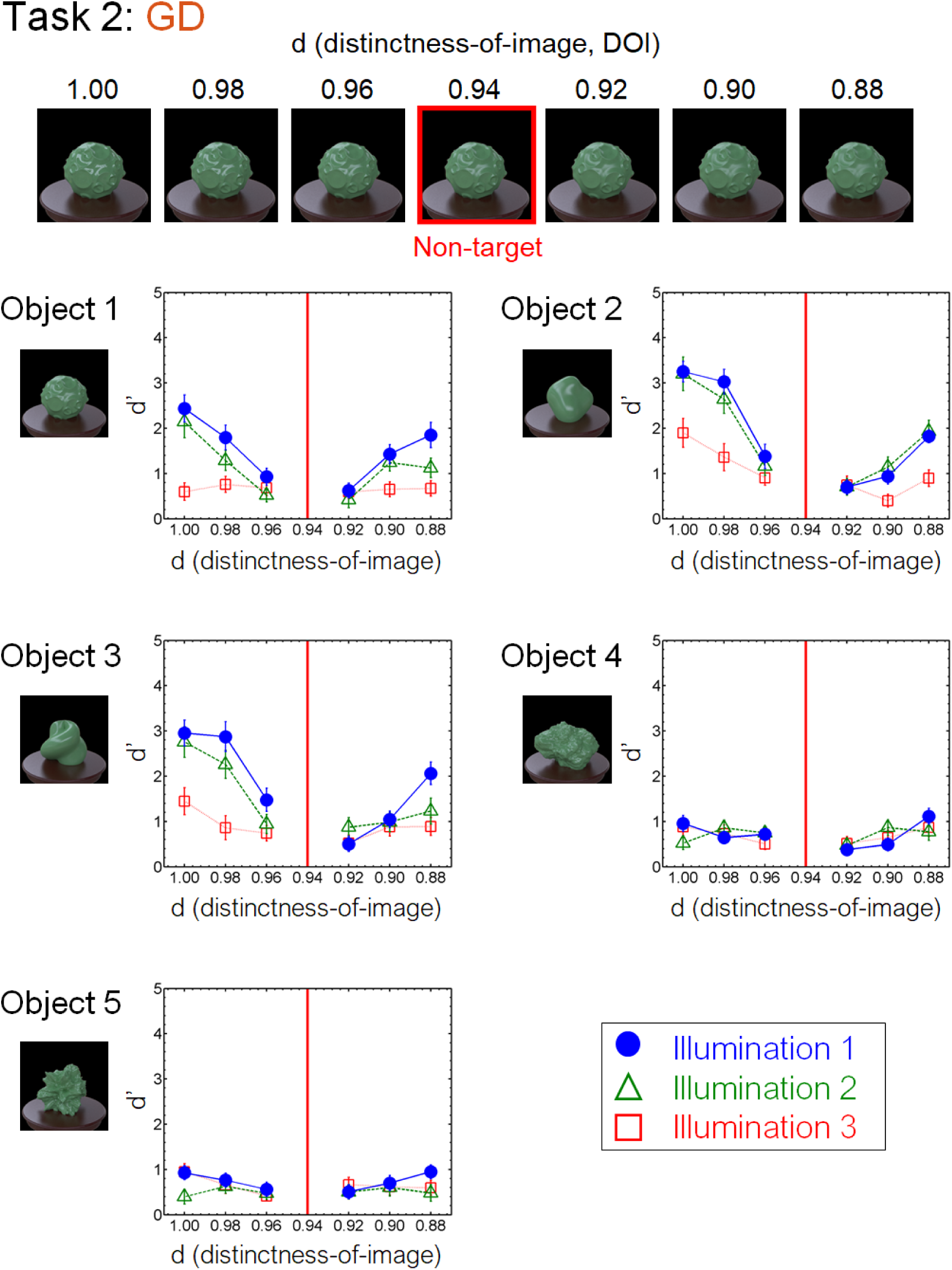
Results of task 2 (GD) in the laboratory experiment.

**Figure 10.**
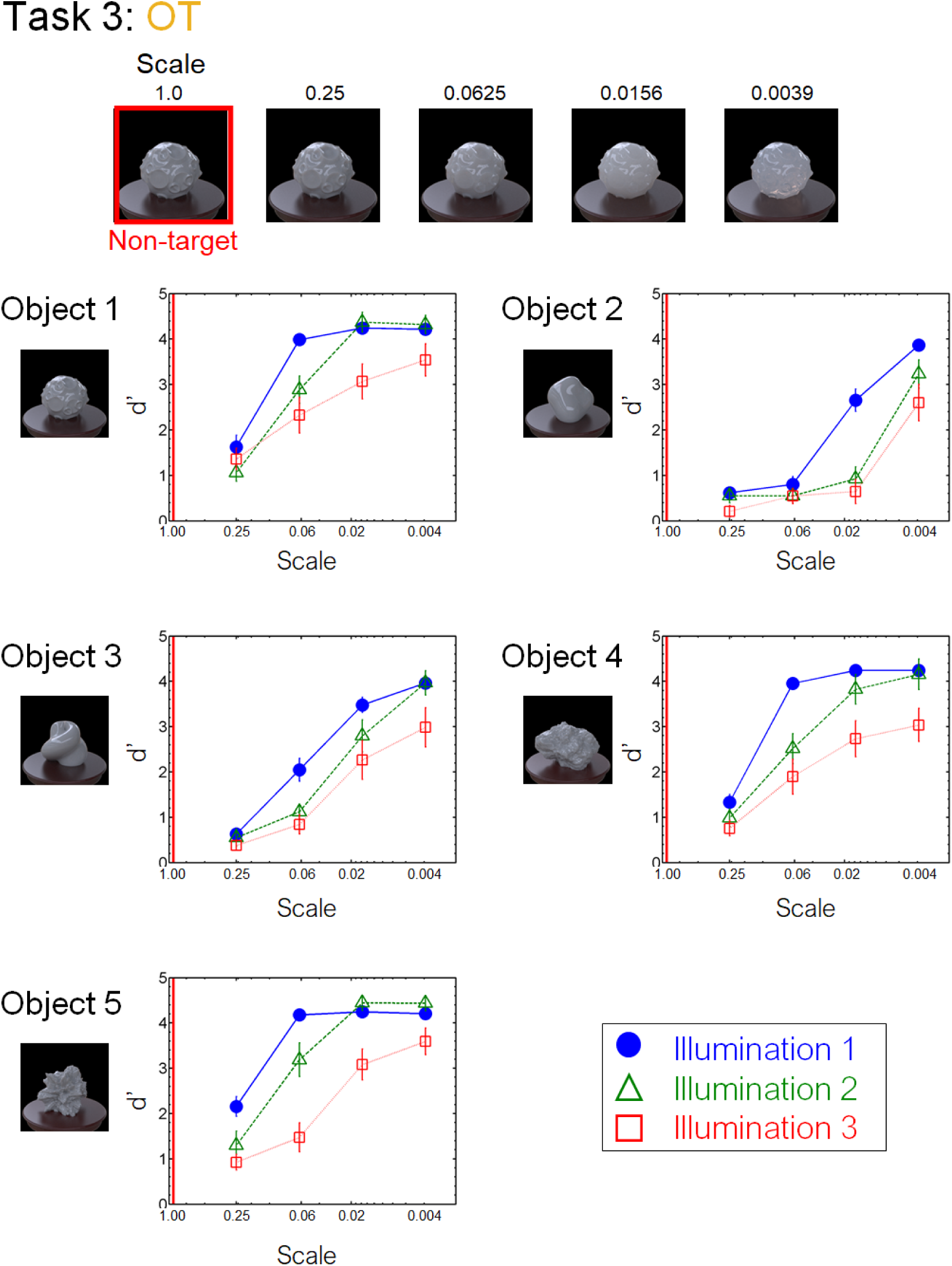
Results of task 3 (OT) in the laboratory experiment.

**Figure 11.**
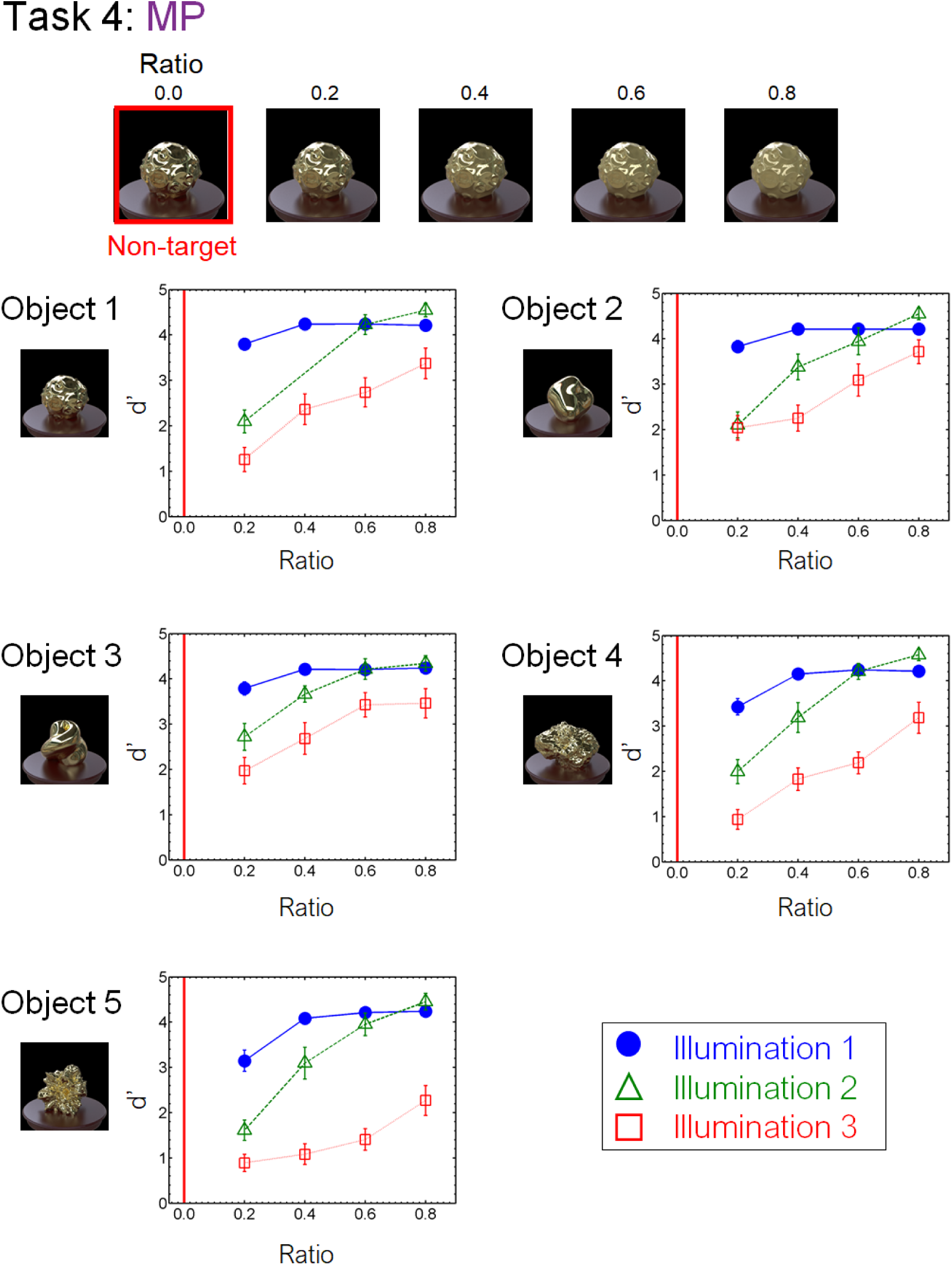
Results of task 4 (MP) in the laboratory experiment. One of the observer data on Object 1 and Illumination 2 is missing due to a mistake in the stimulus presentation.

We also found an illumination dependence in material recognition. We used three illumination conditions, wherein the illumination environments used in a task were identical (Illumination 1), similar to each other (Illumination 2), or largely different from each other (Illumination 3). The results showed that task accuracy decreased as the difference in light probes across the images increased from Illumination 1 to 2 and 3 (Figs. 8-13). This finding not only confirms the large effect of illumination on gloss perception reported before (Fleming et al., 2003; Motoyoshi & Matoba 2012; Zhang et al., 2019), but also demonstrates similarly strong effects of illumination on other material discrimination tasks (OT, MP and MG).

### Intermediate visual feature analysis

One may raise a concern that our observers might make oddity judgments based on differences in low-level superficial image properties such as the object’s mean color. We did not explicitly ask the observers to select one object image in terms of the material appearance. This procedure could lead observers to take a simple strategy unrelated to material judgment. A related question is that, if not such simple properties, are there any intermediate image features in hierarchical visual processing that can explain the observers’ responses? Recent studies have shown that the intermediate processing in the ventral visual stream of humans and monkeys encodes the higher-order image features as computed in texture synthesis algorithms or deep convolutional neural networks (Freeman et al., 2013; Okazawa, Tajima, Komtsu, 2014; 2016, Yamins & Dicarlo, 2015). It has been suggested that the processing in the visual ventral stream also mediates material recognition for static objects (Nishio et al., 2012; 2014, Miyakawa et al., 2017). We asked how such intermediate features possibly processed in material computation can explain the observers’ responses.

More specifically, we analyzed how various image feature differences on each task can explain the observers’ task performance. Each task, i.e., a material dimension with an object under an illumination condition, includes a set of material objects with different combinations of poses (Illumination condition 1) or illuminations (Illumination conditions 2 and 3). These combinations are used as repetition for the behavioral experiment. In the analysis, we chose all combinations for each task and calculated the mean feature distance. We calculated this distance metric using various image features (e.g., pixel statistics or texture statistics) as described below in detail. If the distance metric of each image feature is correlated with human performance, the feature can be diagnostic for human judgments.

We linearly regressed the discrimination sensitivity d’ for each task using the distance metric calculated from various image features. Specifically, we used the texture parameters originally proposed in the literature of texture synthesis by Portilla & Simoncelli (2000). They suggested that natural textures can be synthesized by the probabilistic summary statistics derived from the pixel histogram and the subband distribution, including higher-order statistics such as the correlations across the subband filter outputs. More recently, many studies have shown that the intermediate visual processing in the ventral stream, such as V2 or V4, encodes these texture parameters (Freeman et al., 2011; Okazawa et al., 2013). Following the previous studies (Okazawa et al., 2013), we reduced the original texture parameters by removing redundant features because a large number of parameters make the fitting unreliable. Specifically, we conducted the same reduction as Okazawa et al. (2013), except that 1) we included the mean, sd, and kurtosis of the marginal statistics, as well as the skewness and that 2) we calculated these statistics not only for grayscale images (CIE L* image) but also for color images (CIE a* and CIE b* images). We defined the white XYZ value averaging the diffuse white sphere rendered under each illumination condition and used it to calculate the CIE L*, a*, and b* of each image. We extracted the center 128 x 128 pixels of each image and calculated the texture parameters using the texture synthesis algorithm by Portilla & Simoncelli (1999) with four scales and four orientations. We reduced these original texture parameters of each L*, a*, or b* image to 32 parameters following Okazawa et al. (2013). More details are described in the supplementary tables S1 and S2 of Okazawa et al. (2013). In total, we used 96 parameters for the regression analysis.

We conducted five regressions with different types of parameters to explore the contribution of different statistics. Specifically, we used (1) pixel color means, (2) pixel color statistics, (3) Portilla & Simoncelli’s (PS) grayscale texture statistics, (4) PS grayscale statistics, and pixel color statistics, (5) PS color statistics. The pixel color means and the pixel color statistics were the marginal statistics in the PS texture statistics. The pixel color means indicated the averaged pixel values of each L*a*b* channel. The pixel color statistics indicated the mean, standard deviation, skewness, and kurtosis of each color channel. The number of these parameters was 3 and 12, respectively. For the two conditions, we used a linear regression without regularization to fit the discrimination sensitivity (blue and red in Fig. 14). For the three PS texture statistics conditions (yellow, purple, and green in Fig. 14, respectively), we used the compressed PS statistics as described above. Since the number of parameters for these conditions is large (32, 48, 96, respectively), we used L1-penalized linear least-squares regression (i.e., lasso) to avoid overfitting. We controlled the hyperparameters so that the number of independent variables is 18, where the regression of the PS grayscale statistics condition showed the minimum mean-squared error (MSE).

**Figure 12.**
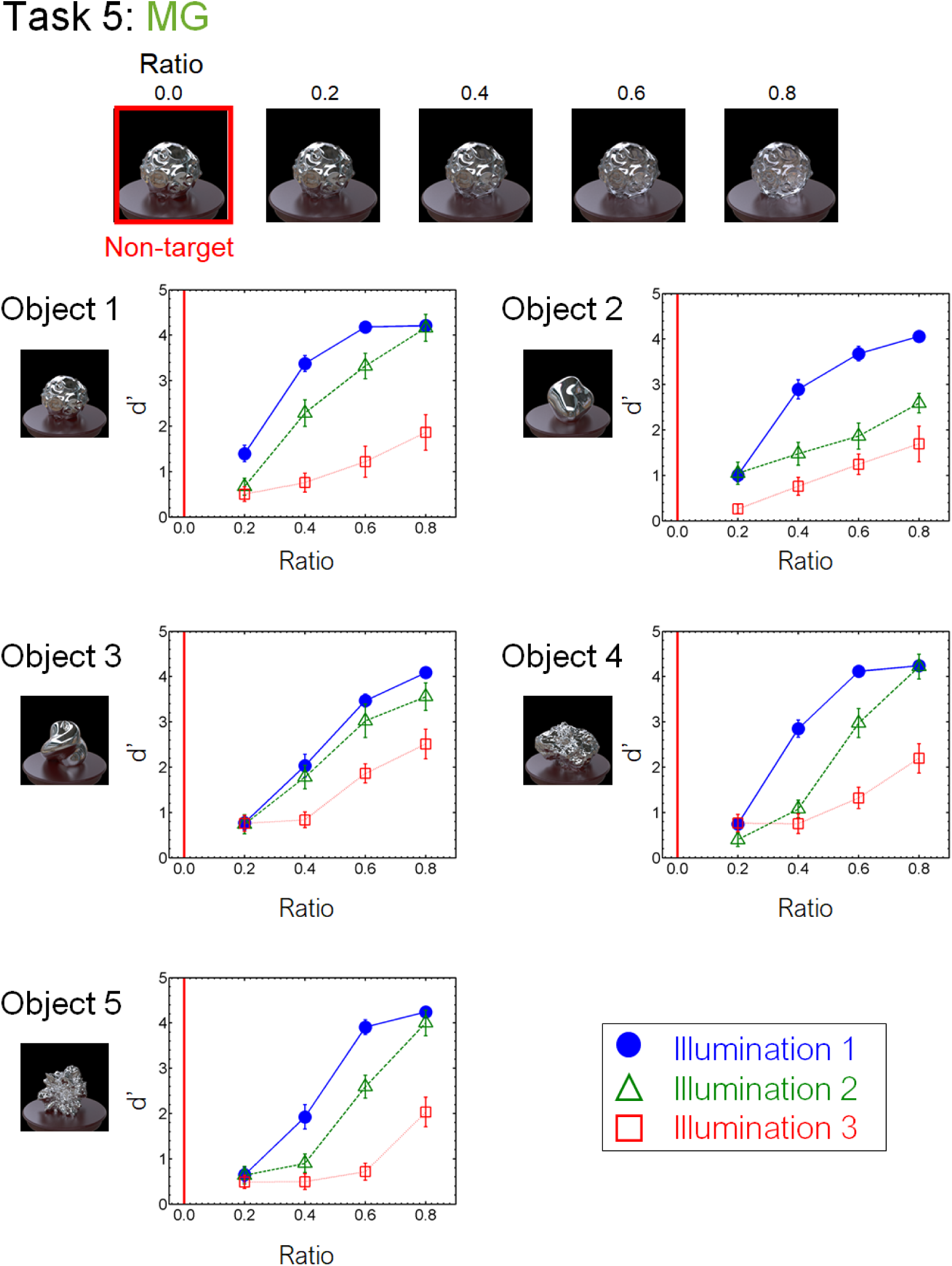
Results of task 5 (MG) in the laboratory experiment.

**Figure 13.**
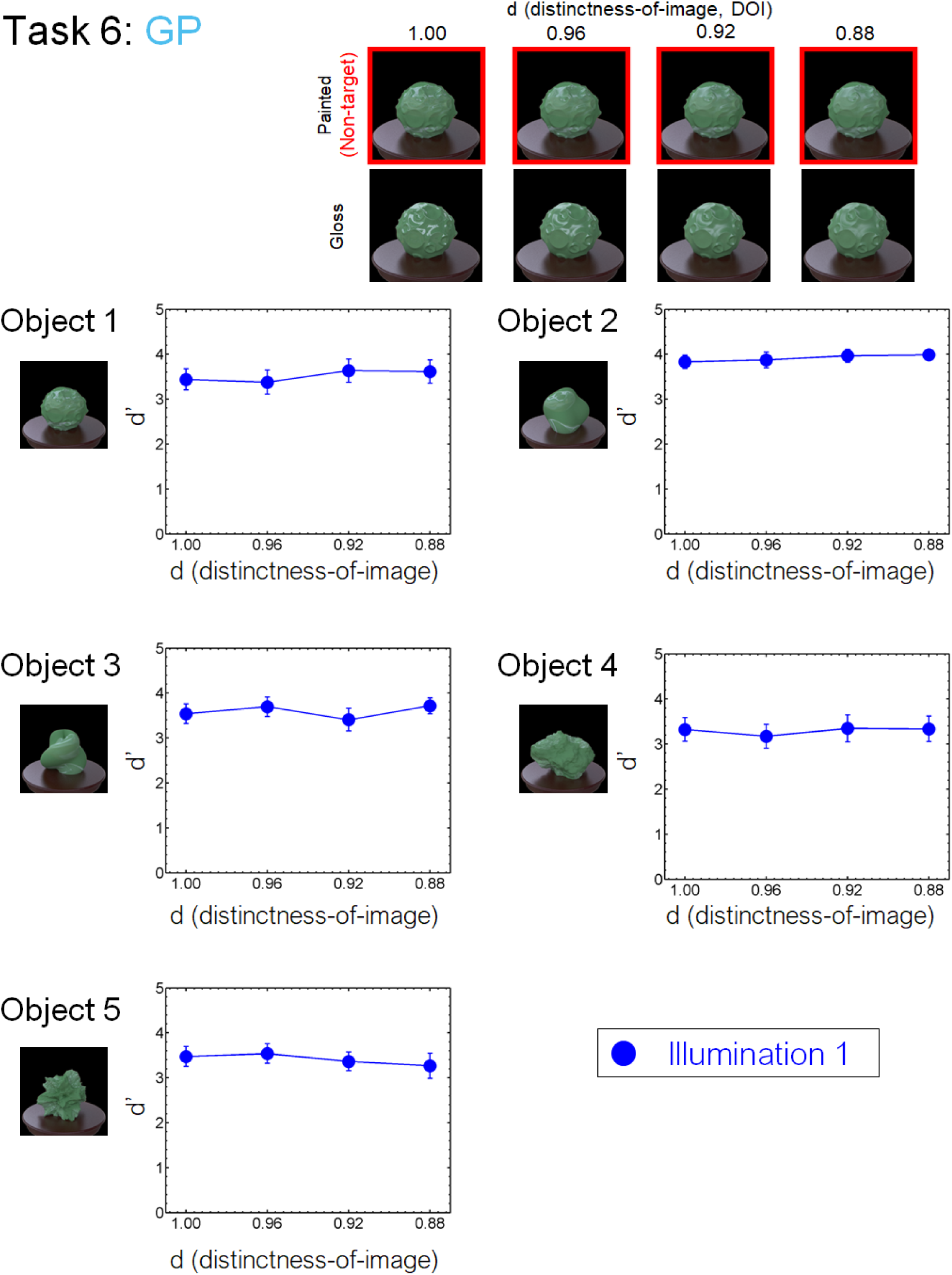
Results of task 6 (GP) in the laboratory experiment.

**Figure 14.**
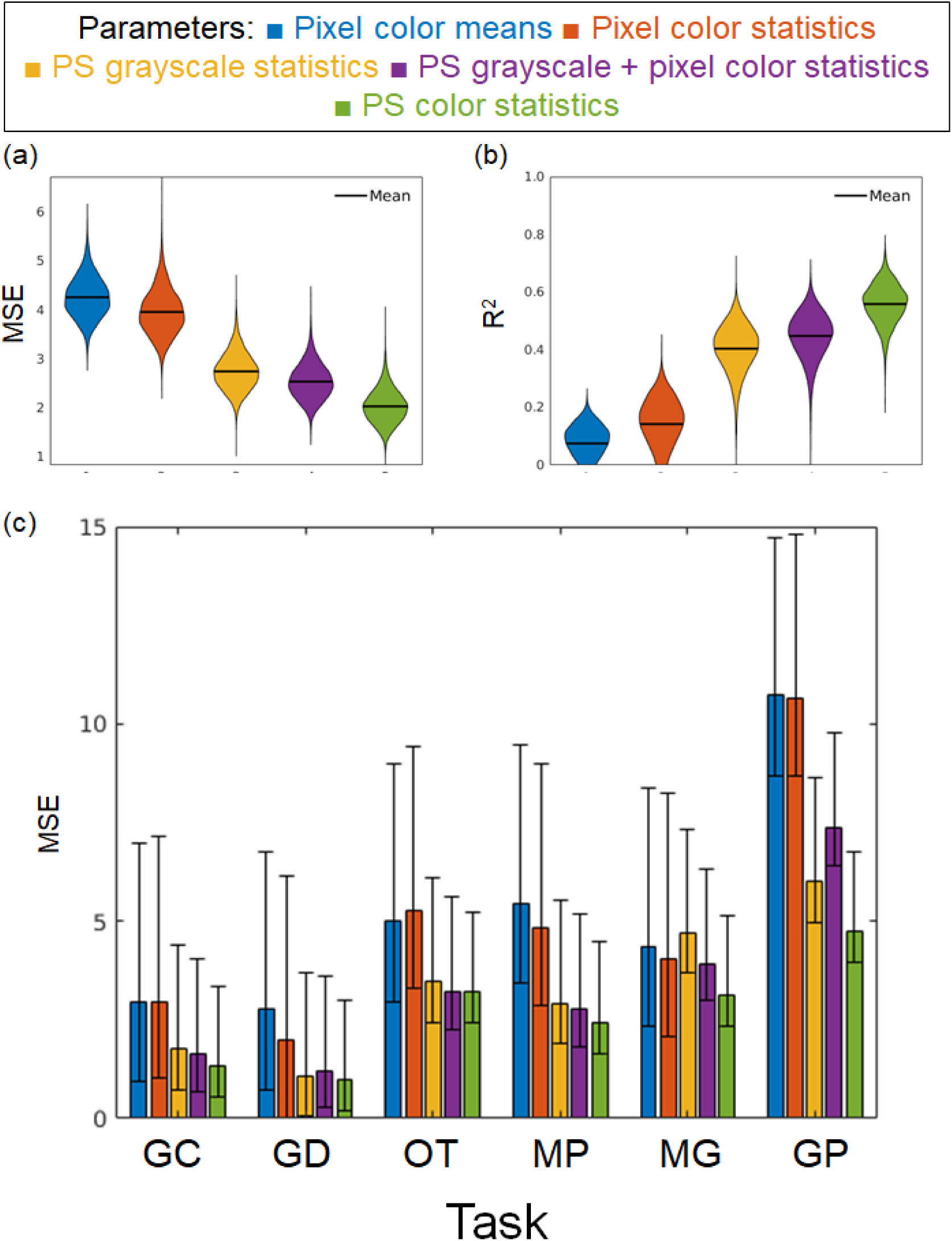
Results of the linear regressions using different parameters. We regressed the human discrimination performance on pixel color means (3 parameters, blue), pixel color statistics (12 parameters, red), Portilla & Simoncelli’s (PS) grayscale texture statistics (regularized 18 parameters, yellow), PS grayscale statistics and pixel color statistics (regularized 18 parameters, purple), or PS color statistics (regularized 18 parameters, purple). (a) Results of the mean squared error (MSE) for each regression. (b) Results of the mean squared error for each regression. These results are shown using a violin plot. (c) Results of the MSEs for each task. The error bars indicate the bootstrap 95% confidence intervals.

We divided all tasks into training and test datasets with a ratio of four to one, respectively, and conducted the above five regressions. The task ratio was kept constant across the training and test datasets. For the training dataset on the lasso regressions, we regressed the discrimination sensitivity using the 5-fold-cross validation. Figure 14 shows the MSE and the determinant coefficient for the test datasets. We resampled the training and test datasets 10000 times and depicted the distribution using a violin plot. First, the predictions based on the color mean statistics didn’t match the observers’ discrimination sensitivity at all (Fig. 14a and 14b). These results suggest that the observers did not simply rely on the mean differences to perform the oddity tasks. The MSE and the determinant coefficient for the marginal statistics condition were more improved when we added the higher-order statistics (marginal statistics condition, PS grayscale statistics condition, and PS color statistics condition). Since the regularization parameter is controlled under the PS color and grayscale statistics conditions, these results cannot be ascribed to the number of independent variables. It is noteworthy that even when all the PS color statistics are used, the prediction is not sufficient to explain observers’ discrimination performance. This finding suggests that human material judgments also rely on higher-order features the PS statistics do not cover. One possible future direction is to use the intermediate activation of the deep neural networks. To support this direction, we include in our database the activation data of VGG-19, a feedforward convolutional neural network, for our image dataset and the analysis about how the dataset is represented in each layer (Appendix C). In short, our dataset images were clustered in higher layers of the pretrained network according to object differences, and the material differences were represented in each object cluster.

### Individual differences

Next, we evaluated the individual differences of each task in the Japanese adult population. Figure 15 shows the histogram of the response accuracy for each observer in the crowdsourcing and laboratory experiments. For the crowdsourcing experiment, the number of observers of illumination conditions 1, 2, and 3 was 416, 411, and 405, respectively. For the laboratory experiment, the number of observers was 20. For each condition, the probability of a correct response was calculated by averaging the responses of each observer across objects and task difficulties. The standard deviations of tasks 1 to 6 under illumination condition 1 are .14, .11, .12, .12, .12, and .23, indicating a particularly large individual difference for task 6 (GP). The standard deviation under illumination conditions 2 and 3 ranged from .09 to .18. It should be also noted that most of the conditions show unimodal distributions, while task 6 (GP) shows a nearly uniform distribution. This finding suggests that individual differences in discrimination ability of the spatial consistency of specular highlights are larger than those for other material properties, including glossiness contrast and distinctness-of-image (GC, and GD).

**Figure 15.**
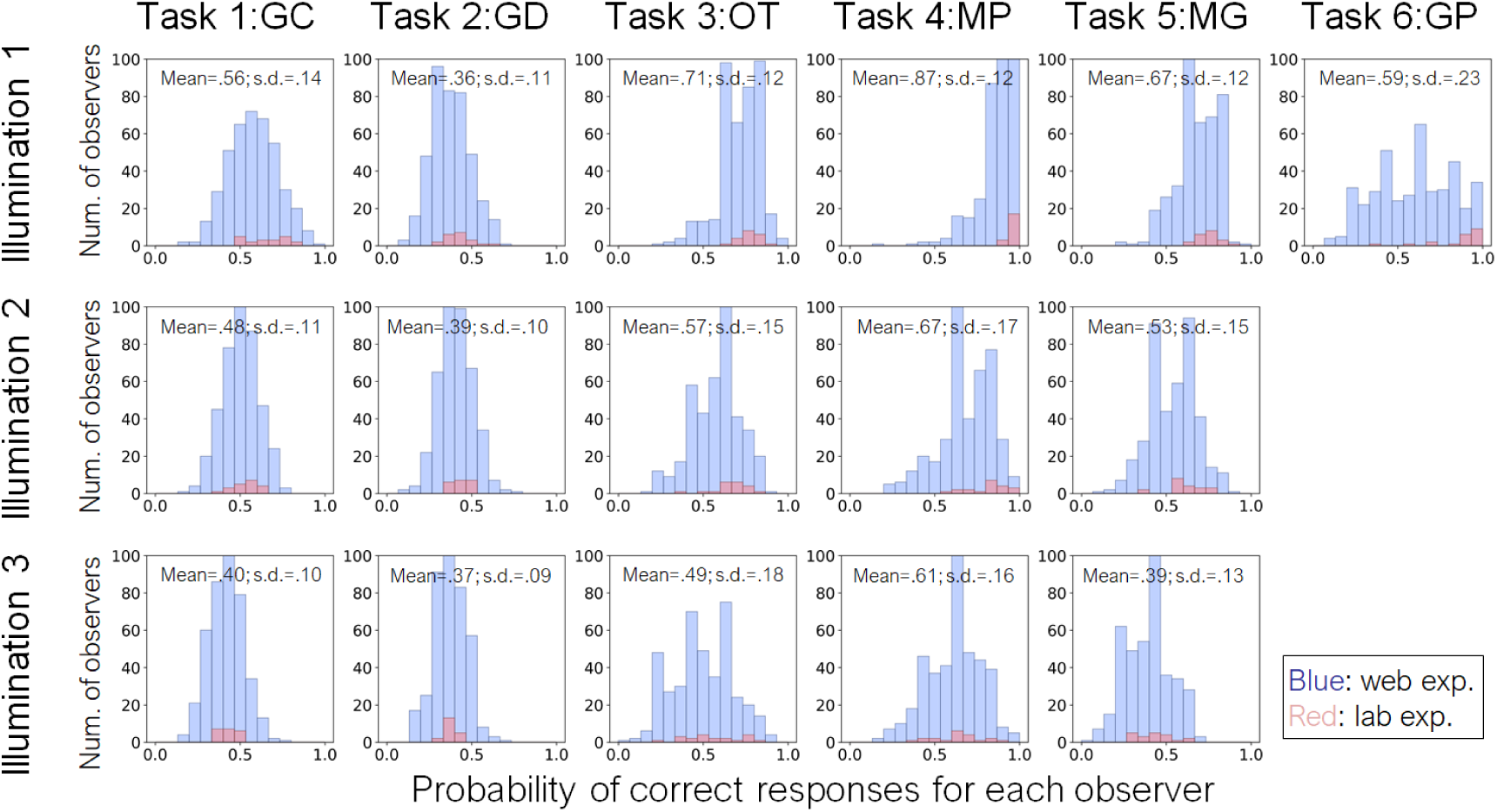
Histogram of response accuracy for each observer in the crowdsourcing (blue) and lab (red) experiments. Different panels indicate different material tasks and illumination conditions. For each condition, the probability of a correct response was calculated by averaging the responses of each observer across objects and task difficulties. The histograms of crowdsourcing and lab experiments are overlayed in each panel. The mean and standard deviation of each distribution are shown in each panel.

## Discussion

The present study aimed to construct a database of material images annotated with the results of human discrimination tasks. We created material images that varied in six different material dimensions on the basis of the previous material-recognition studies. Our dataset includes various objects and illuminations so that users can comprehensively investigate the effects of these physical causes on material recognition. The results of psychophysical experiments showed that most of the task difficulty could be appropriately controlled by manipulating the material parameters. Furthermore, analysis of visual feature showed that the parameters of higher-order color texture statistics (Fig. 14, PS color statistics) can partially, but not completely, explain task performance. One crucial point of our dataset is that we used a non-verbal procedure to collect the observers’ data. Since this procedure is widely used in babies, brain-injured participants, and animals, the current behavioral data can be a benchmark for more diverse research fields.

Since we comprehensively investigated the material recognition using a structured dataset, our dataset itself revealed novel findings about material recognition. For instance, the present results showed that the performance of the tasks in the crowdsourcing experiment was strongly correlated with that in the laboratory experiment. This suggests that the dataset has enough tolerance to conduct new experiments involving a variety of observers and experimental conditions. Another is that geometry dependency on material recognition emerges similarly in different material attributes such as gloss distinctness-of-image or translucency (Fig. 10). Specifically, the translucency discrimination sensitivity was high when the object had rugged surfaces (e.g., Object 1, 4, & 5). Some studies have shown that physically prominent features of translucent objects appear around sharp corners on the surface (Fleming et al., 2005; Gkioulekas et al., 2013). One possibility is that the diagnostic features for translucent perception lie in the edge/corner of a translucent object and our rugged objects included much information to judge translucency. More recently, Xiao et al. (2020) investigated the effect of geometry on translucency perception. In their experiments, they changed the smoothness of the object edges. In agreement with our findings, the edge modulation was critical to the translucency perception. Specifically, the object with the smooth edge was perceived as more translucent than the sharp one.

Another finding is that the ability to discriminate the spatial consistency of specular highlights in glossiness perception has large individual differences, although other glossiness discrimination tasks do not show such large differences. Some studies suggest that image statistics are diagnostic for glossiness perception (Adelson, 2001; Motoyoshi et al., 2008). However, when specular highlights of an object image are inconsistent in terms of their position and/or orientation with respect to the diffuse shading component, they look more like white blobs produced by surface reflectance changes (Beck & Prazdny, 1981; Kim et al., 2011; Marlow et al., 2011). This is why the highlight-inconsistency effect is considered to be a counterexample to the image statistics explanation. The large individual differences suggest that the discrimination of the spatial consistency of specular highlights may be mediated by a different, and possibly more complicated, mechanism than that responsible the glossiness contrast/distinctness-of-image discrimination. In agreement with this notion, Sawayama and Nishida (2018) showed that highlight inconsistency is discriminated by different image gradient features from those used in the human material computation. This suggests that the glossiness computation is mediated by multiple stages, i.e., one is to discriminate different materials on a surface for extracting a region-of-interest (ROI), and another is to compute the degree of glossiness in the ROI as shown in Motoyoshi et al. (2007).

One may have a concern that the intermediate objects in tasks 4 and 5 are physically infeasible because they are a mixture of two physically distinct materials. However, our stimuli do not look so unrealistic. The dielectric/metal materials are distinct material categories when considering an object with a uniform single material, but many daily objects surrounding us however are a mixture of various materials, and we often see a plastic object coated by a metallic material. We can regard our intermediate materials as an approximation of such coated materials. In addition, continuously connecting distinct categories is common in various research fields such as speech recognition (e.g., Grey & Gordon, 1978) or face recognition (e.g., Turk et al., 2002), especially to elucidate what stimulus image features are involved in the processing. Considering the literature, we think our intermediate approach is reasonable.

Although our database includes diverse material dimensions, they are still not enough to cover the full range of natural materials. One example is cloth (Xiao et a., 2016; Bi & Xiao, 2016; Bi et al., 2018; 2019). Cloth materials are ubiquitous in everyday environments. A reason we did not include this class of materials is that it has been shown that the cloth perception strongly relies on dynamic information (Bi et al., 2018; 2019). Because of the limited experimental time, our database currently focuses on static images. This is why other materials related to dynamic information (reviewed by Nishida et al., 2018) related to the perception of liquidness (Kawabe et al., 2015), viscosity (Kawabe et al., 2015, van Assen & Fleming, 2018), stiffness (Paulun et al., 2017; Schmid & Doerschner, 2018), etc., were not used in the current investigation. In addition, the perception of wetness (Sawayama, Adelson, & Nishida, 2017) and the fineness of surface microstructures (Sawayama, Nishida, & Shinya, 2017) were not investigated because of the difficulty of continuously controlling physical material parameters by using identical geometries of other tasks. Since we only used five geometries, material perceptions derived from object mechanical properties were not investigated either (Schmidt et al., 2017). A crucial point is that we share our source code to reproduce images. We hope to remove obstacles to constructing a new dataset and contribute to future work on material recognition. Sharing the datasets with the source code should make researchers easily conduct a new experiment in this literature. For instance, we measured the discrimination sensitivities in our experiments from one side of the materials in tasks 3, 4, and 5 (i.e., opaque, gold, and silver). The sensitivities from the other side (i.e., transparent, plastic, and glass) could be slightly different from the current results. Researchers can easily render new images of different material parameters in the same scene condition and conduct a new investigation.

Our datasets also highlighted the difficulty of choosing appropriate parameters that cover the full range of the material sensitivity. We chose the stimulus parameters based on the preliminary experiments. We tried to choose the parameters so that we can measure the sensitivity of each task in the full range, i.e., from the chance level to the maximum accuracy. However, we found large individual differences in some tasks, e.g., task 6, and they resulted in the partial measurement of the narrow sensitivity range. This unpredictability is one of the difficulties of producing the large size of the dataset. The current findings should contribute to the future attempt making material image datasets.

Our dataset focuses on expanding the previous findings as to material recognition into more diverse research fields. From the view of a global standard dataset, our dataset has several limitations as described above. However, it did contribute to this expansion purpose. Specifically, several research groups of behavioral science, computer science, and neuroscience have on-going projects utilizing our dataset, and some findings have already been reported at conferences and journals. Kawasaki et al. (2019) used our dataset to explore the role of the monkey ITC on material perception by using the electrocorticography (ECoG) recordings. Tsuda et al. (2020) investigated the role of working memory on material processing using our dataset. Koumura et al. (2018) explored how mid-level features in deep convolutional neural networks can explain human behavioral data. Imura et al. (2017) compared the discrimination performance of children and adults. The attention and memory roles in material recognition are also investigated by Takakura et al. (2017).

## Conclusion

We constructed image and observer database for material recognition experiments. We collected observation data about material discrimination in tasks that had a non-verbal procedure for six material dimensions and several task difficulties. The results of psychophysical experiments in laboratory and crowdsourcing environments showed that the performance of the tasks in the crowdsourcing experiment was strongly correlated with the performance of the tasks in the laboratory experiment. In addition, by using the above comprehensive data, we showed novel findings on the perception of translucence and glossiness. Not only can the database be used as benchmark data for neuroscience and psychophysics studies on the material recognition capability of healthy adult humans; it can also be used in cross-cultural, cross-species, brain-dysfunction, and developmental studies of humans and animals.

## Acknowledgements

This work was supported by a Grant-in-Aid for Scientific Research on Innovative Areas from the Japan Society of Promotion of Science to SN and YD (JSPS KAKENHI Grant Numbers JP15H05915, JP15H05924, JP20H05954, JP20H05957).

Commercial relationships: none.

## Competing interests

The authors declare no competing financial interests.

## Appendix A

The results of the crowdsourcing experiment are shown in Figures A1 to A6. The same experiments were also conducted in the laboratory environment, and their results are shown in Figures 8 to 13.

**Figure A1.**
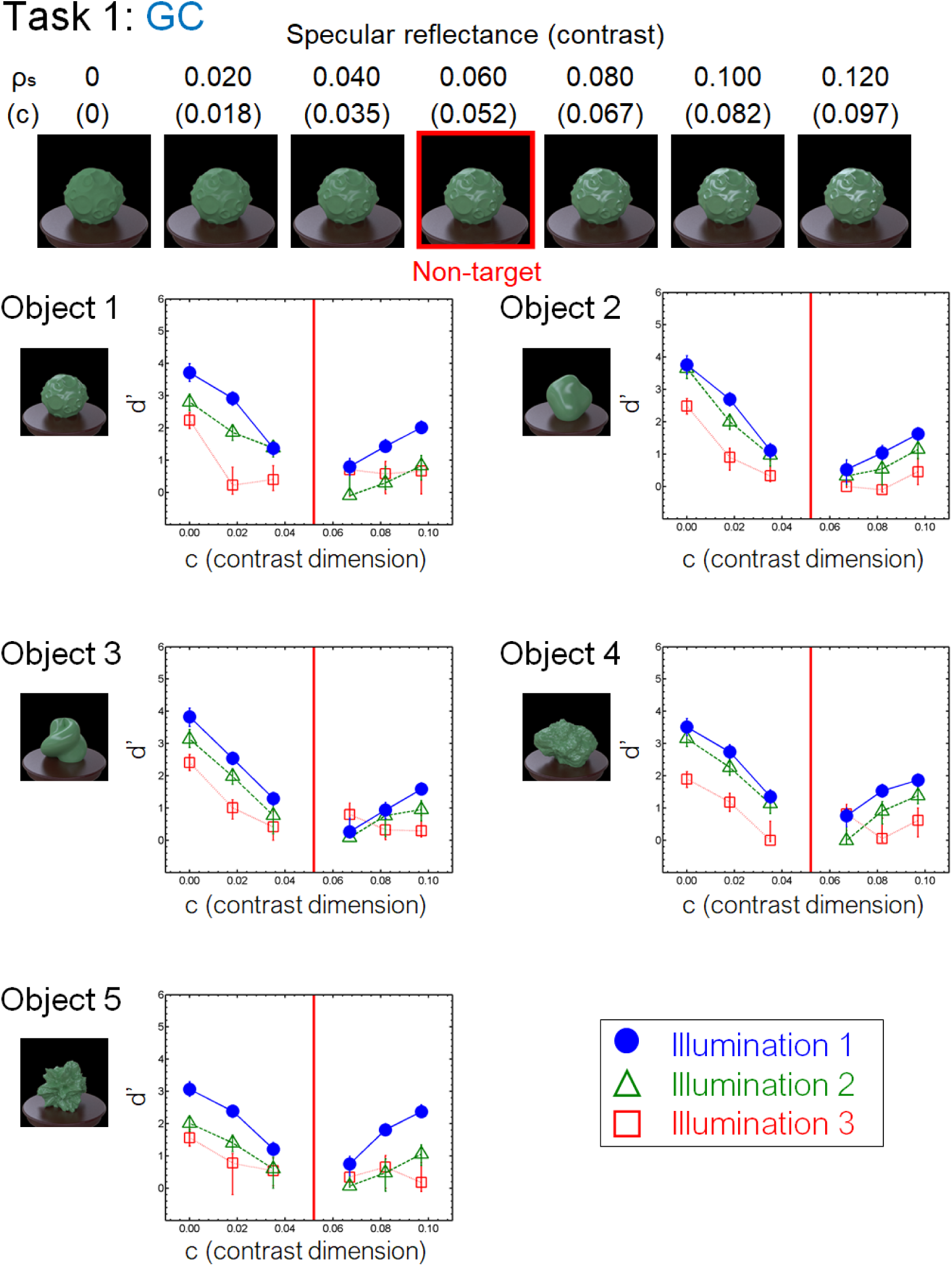
Results of task 1 (GC) in the crowdsourcing experiment. Different panels show different objects. Different stmbols in each panel depict different illumination conditions. The vertical red line in each panel indicates the parameter of the non-target stimulus. Error bars indicate the 95% bootstrap confidence intervals.

**Figure A2.**
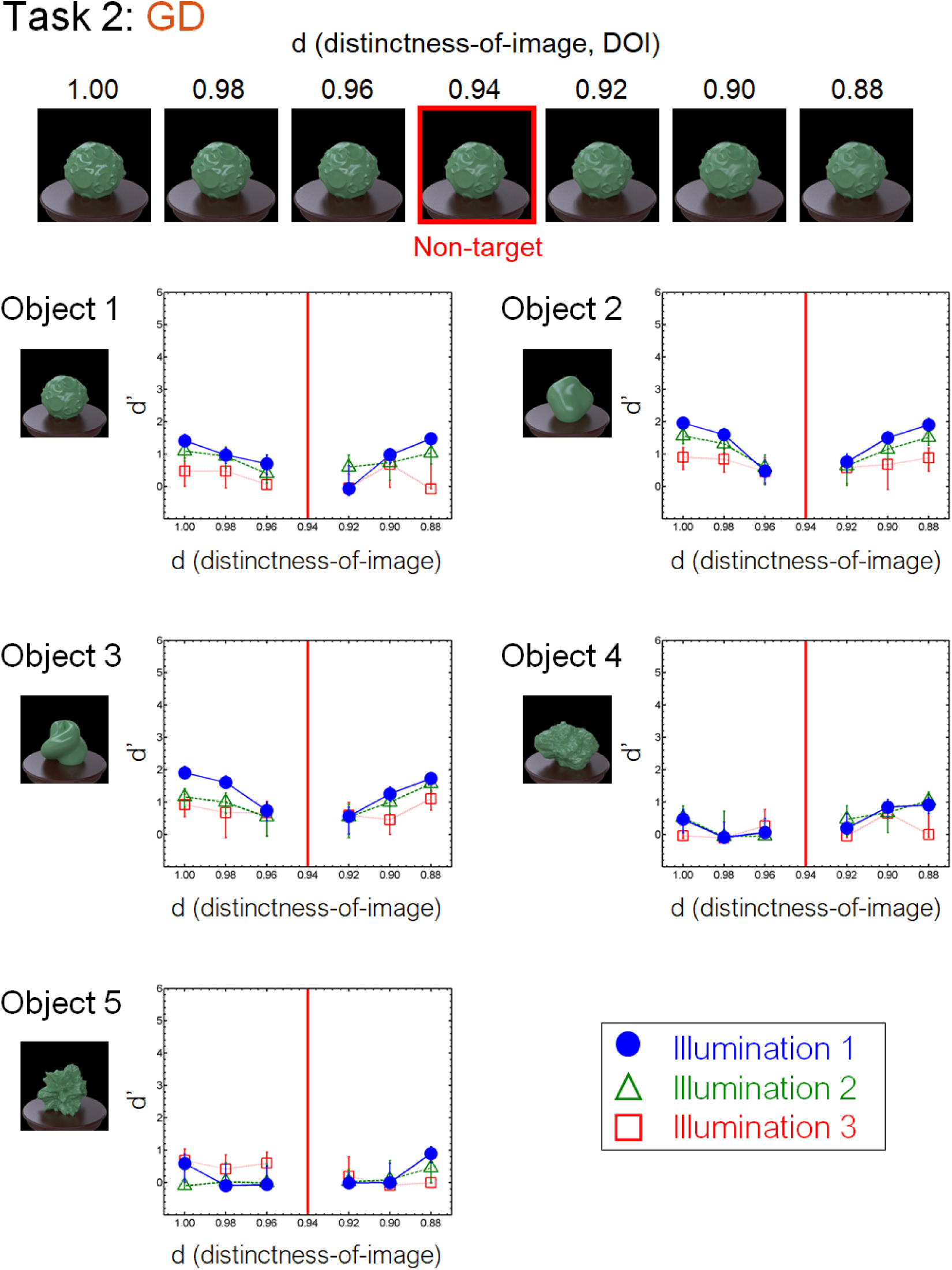
Results of task 2 (GD) in the crowdsourcing experiment.

**Figure A3.**
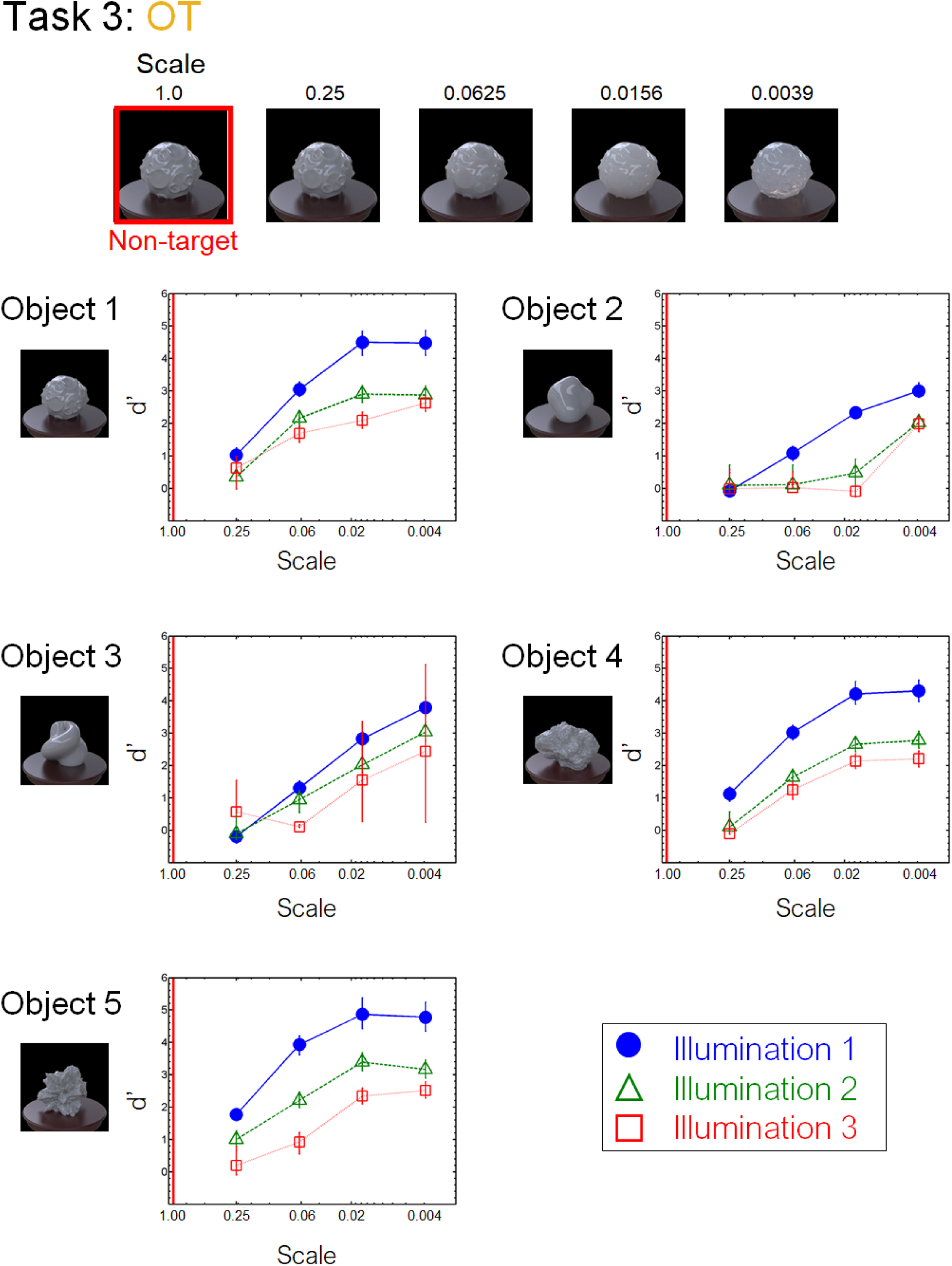
Results of task 3 (OT) in the crowdsourcing experiment.

**Figure A4.**
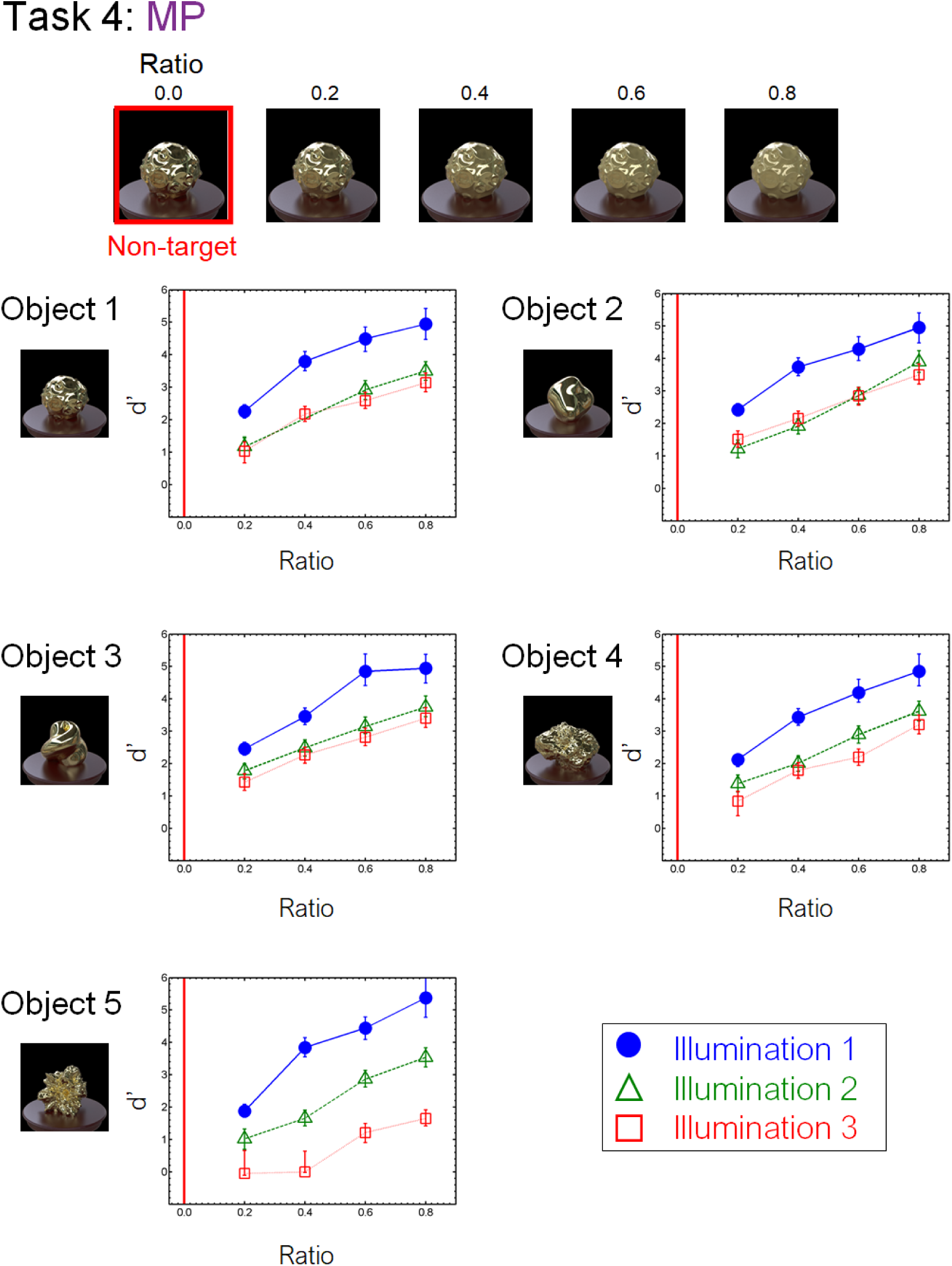
Results of task 4 (MP) in the crowdsourcing experiment.

**Figure A5.**
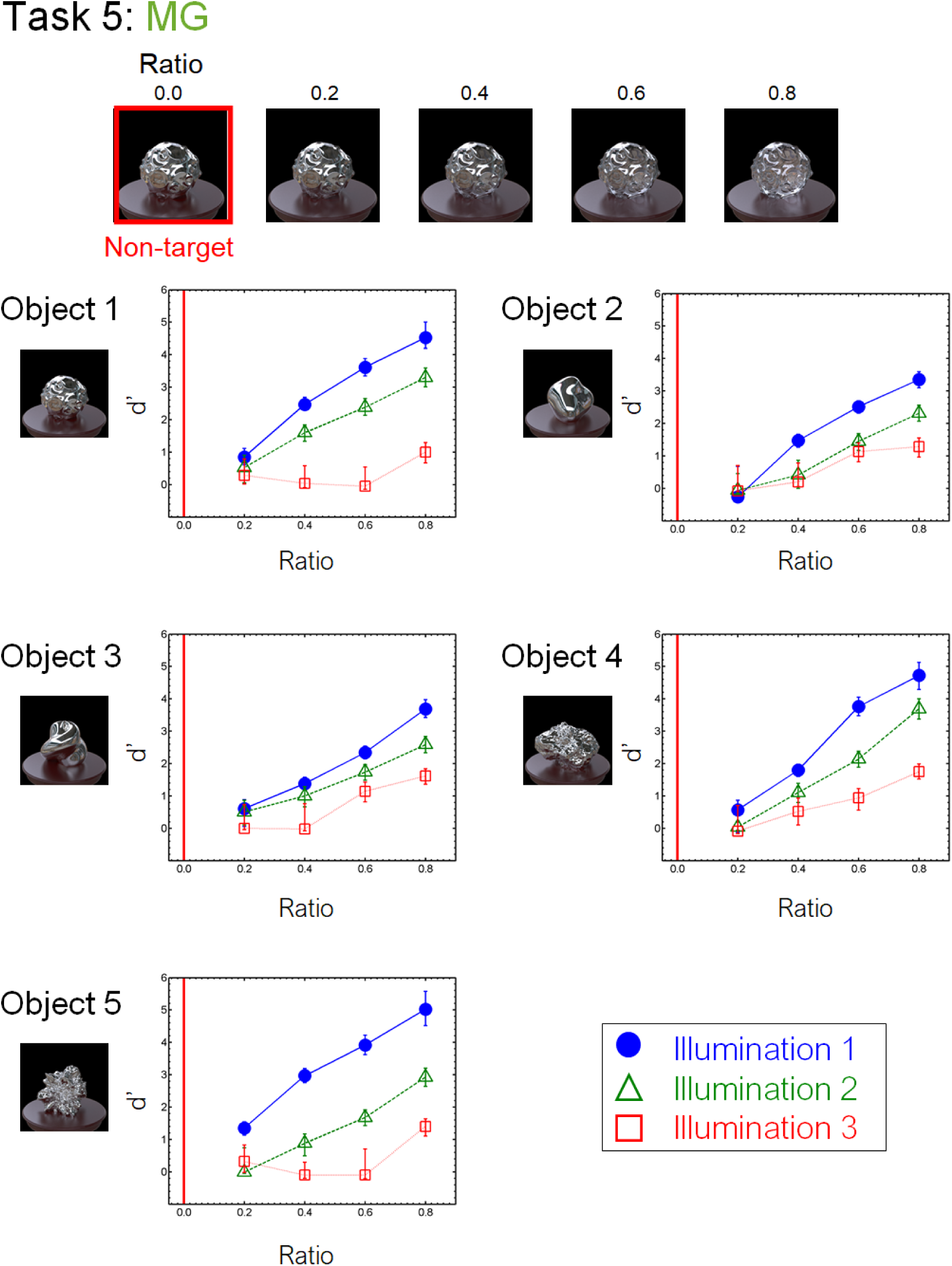
Results of task 5 (MG) in the crowdsourcing experiment.

**Figure A6.**
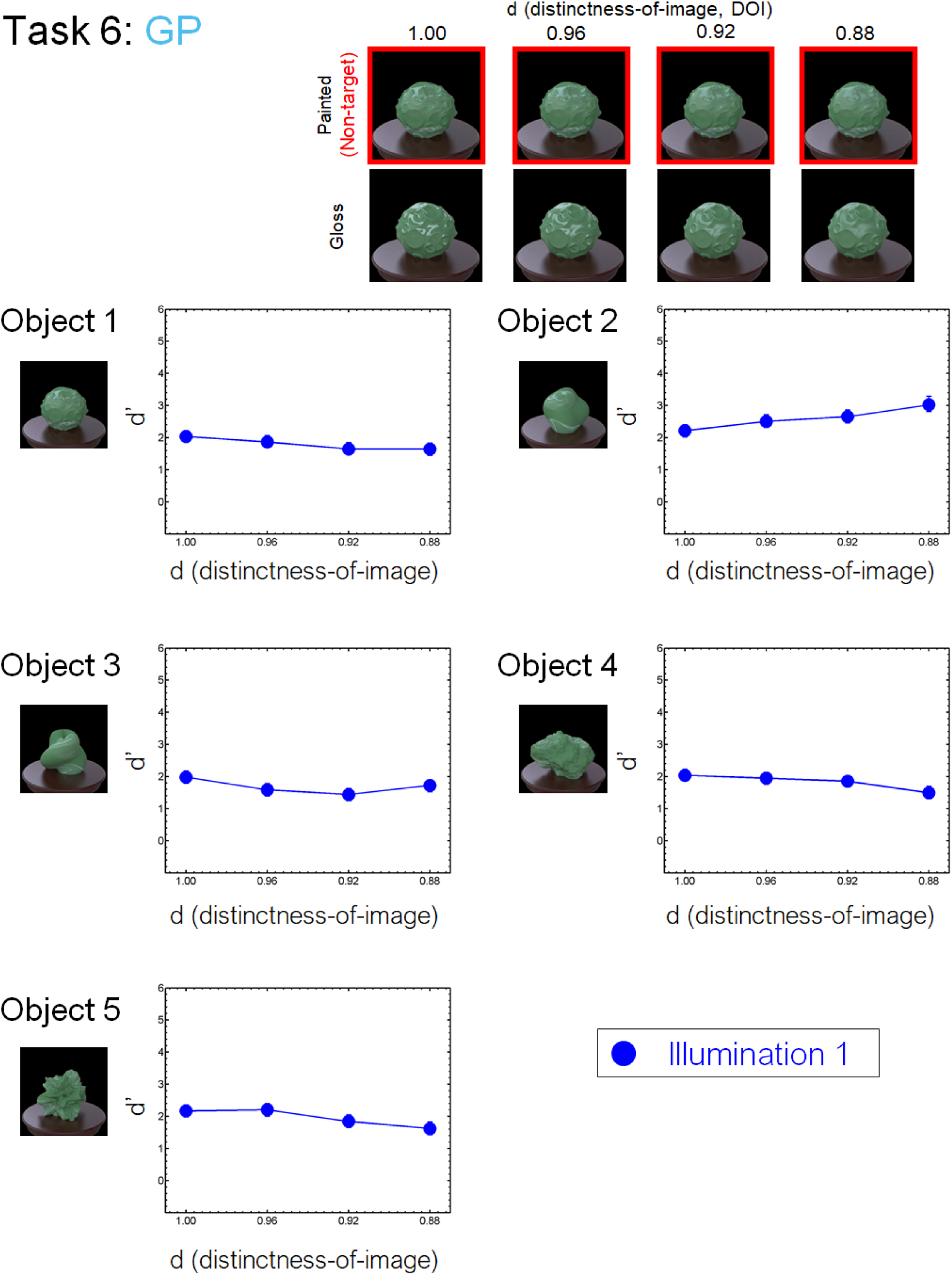
Results of task 6 (GP) in the crowdsourcing experiment.

## Appendix B

### Data records

The database is available at https://github.com/mswym/material_dataset. Figure B1 shows the data structure. The standard data are divided into three folders according to the illumination conditions. Each illumination condition folder contains folders of the material tasks (Task 1 to 6). Each material task folder includes experimental task folders. Each experimental task folder corresponds to one task in the behavioural experiments. The name of each folder indicates the illumination condition, object, material task, and task level. For instance, the name “Il1_obj1_Task1_06_12” indicates illumination condition 1 (i.e., Il1), object 1 (i.e., obj1), task 1 (Task1), contrast of 0.06 for the non-target stimulus, and contrast of 0.12 for the comparison stimulus.

Each task folder contains the two folders named “1” and “0”. The images in the folder “0” indicate the non-target stimuli, while the images in the folder “1” are the target stimuli. Under illumination condition 1, three images are randomly selected from folder “0”, and one correct image is selected from folder “1”. Five images with different poses are stored in each “1” or “0” folder for illumination condition 1, while three images with different illuminations are stored for illumination conditions 2 and 3. The images in the database are in .png format and have a size of 512 x 512 px. In addition, standard observer data are placed on the top layer in the database in a .csv file. The file includes observer data including the probability of the correct response and the sensitivity d’ for each task in the crowdsourcing and laboratory experiments.

**Figure B1.**
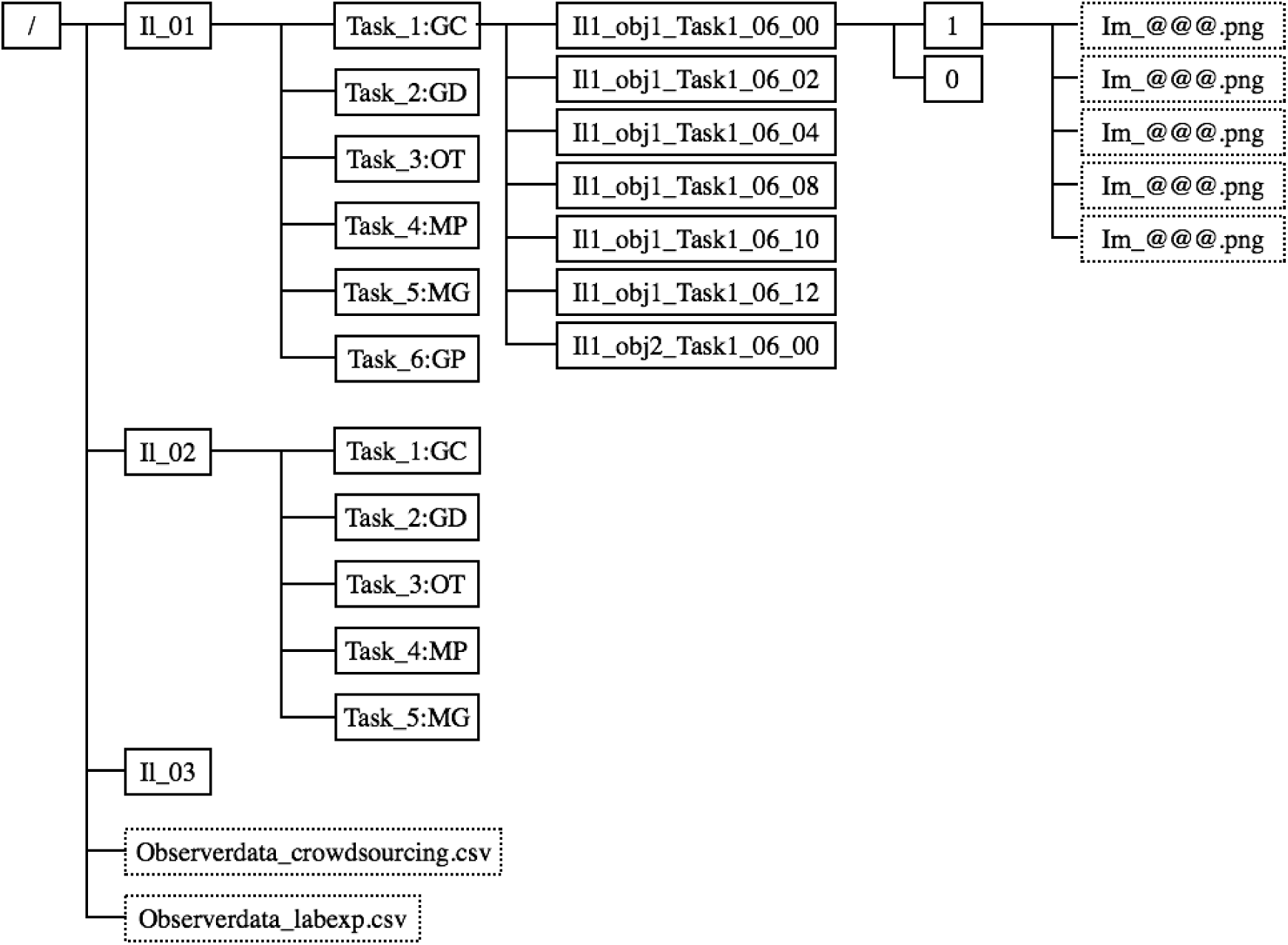
Data structure in the database. Solid rectangles indicate a folder, while the dashed ones indicate a file.

## Appendix C

We analyzed how our datasets are represented in convolutional neural networks (CNNs). We extracted the visual features from each intermediate layer of a CNN. We used the VGGNet16 (Simonyan and Zisserman, 2014), pre-trained for the object recognition task using ImageNet 2012 (Russakovsky et al., 2015), and computed the activation of thirty convolution layers and three fully-connected-layers of the model. To reduce the number of dimensions, we spatially averaged each channel’s activation. Thus, we obtained the multidimensional activation vector for each layer with the dimension number of the channels.

Figures C1 to C4 show the t-SNE embedding of each layer (Maaten & Hinton, 2008). Figure C1 shows the results of the first convolution layer (conv 1_1), the last convolution layer (conv 5_3), and the third fully-connected-layer. Each plot indicates each material image. Different panels in each column mean different labelings based on task, object, and illumination, as shown in the legends. Figure C2 shows the embeddings of all the layers, which are colored by different tasks. Figures C3 and C4 show the same embeddings as Figure C2, except colored according to different objects and illuminations, respectively.

The embedding of the first convolution layer (conv 1_1) showed the clusters according to task differences, especially MG, MP, and OT clusters. In contrast, this embedding didn’t show any object-based clusters. Earlier layers are generally sensitive to lower image features. Different tasks have different colors in our datasets, except that the tasks GC, GD, GP share similar green colors. In addition, some clusters of illumination condition 3 emerged in the first layer embedding. The pixel color distribution of illumination condition 3 is also largely different from the others. These results suggest that the first layer code such lower image features.

The embeddings of the last convolution layer and the third fully-connected layer showed the clusters according to object differences. Different tasks and illuminations are separately distributed within each object cluster. Although the embedding is clustered according to object differences, it didn’t show the separation between Objects 2 and 3. This finding is consistent with human discrimination performance. The results of behavioral experiments showed that the task accuracies of Objects 2 and 3 were similar to each other and different from other object conditions, especially on Task GD and OT.

**Figure C1.**
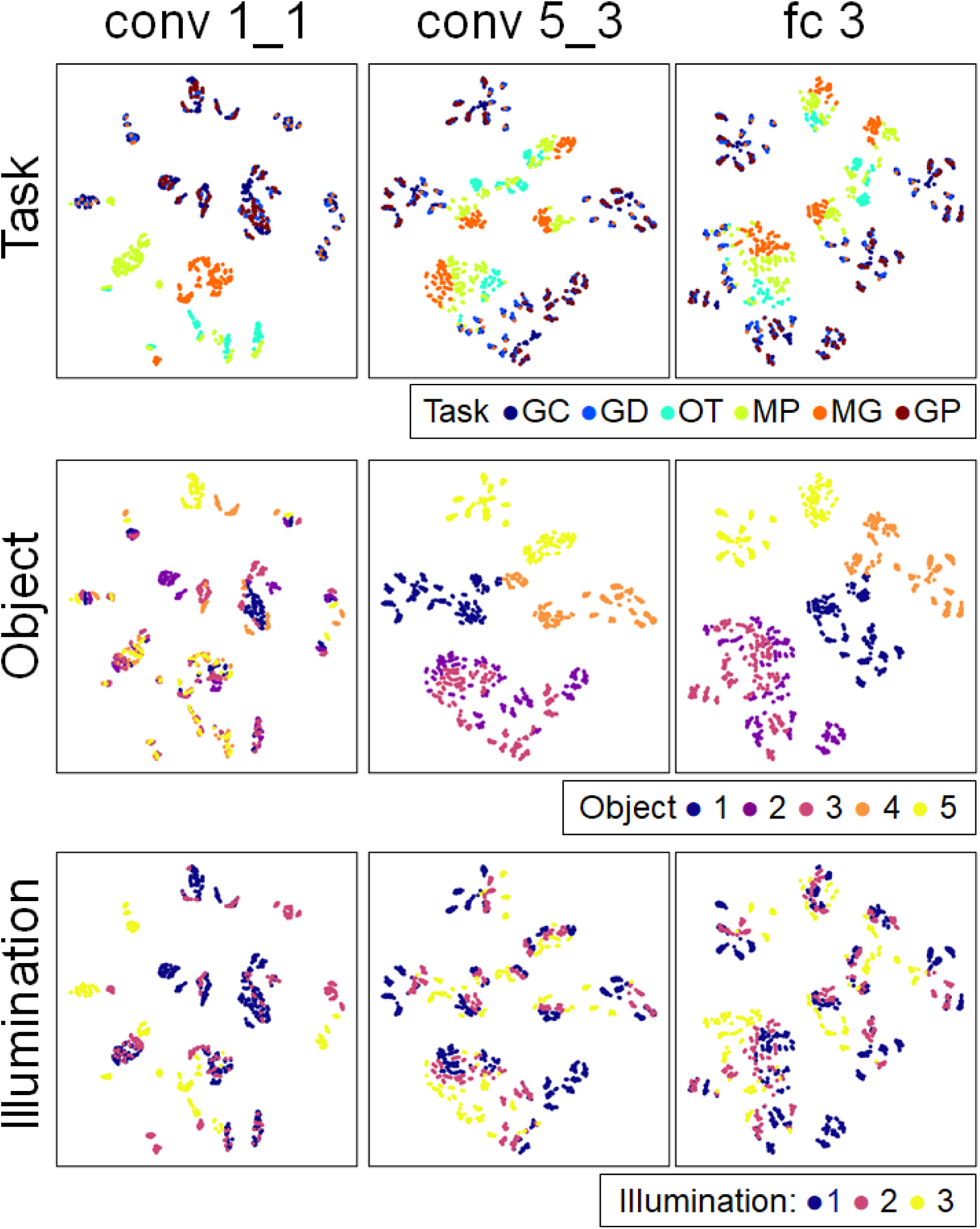
Embedding spaces of intermediate features of a deep neural network trained for object recognition. The top, center, and bottom rows show the same embedding spaces with different color symbols as shown in the legend. The left, middle, and right columns are the results of the first convolution layer (conv1_1), the final convolution layer (conv5_3), and the third fully connected layer (fc 3), respectively.

**Figure C2.**
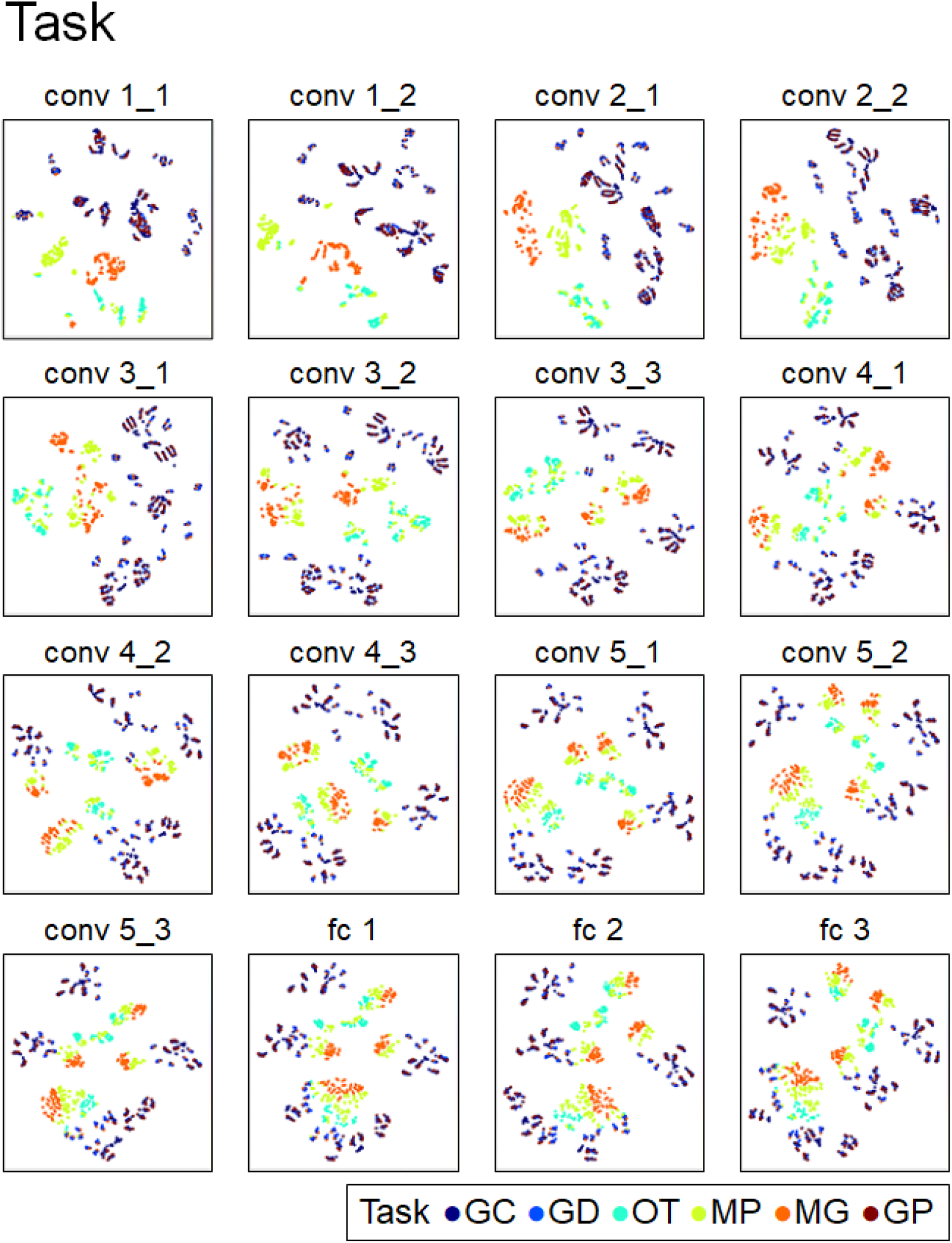
Embedding spaces of intermediate features of a deep neural network trained for object recognition. Results of all the 16 layers are shown with coloring different tasks.

**Figure C3.**
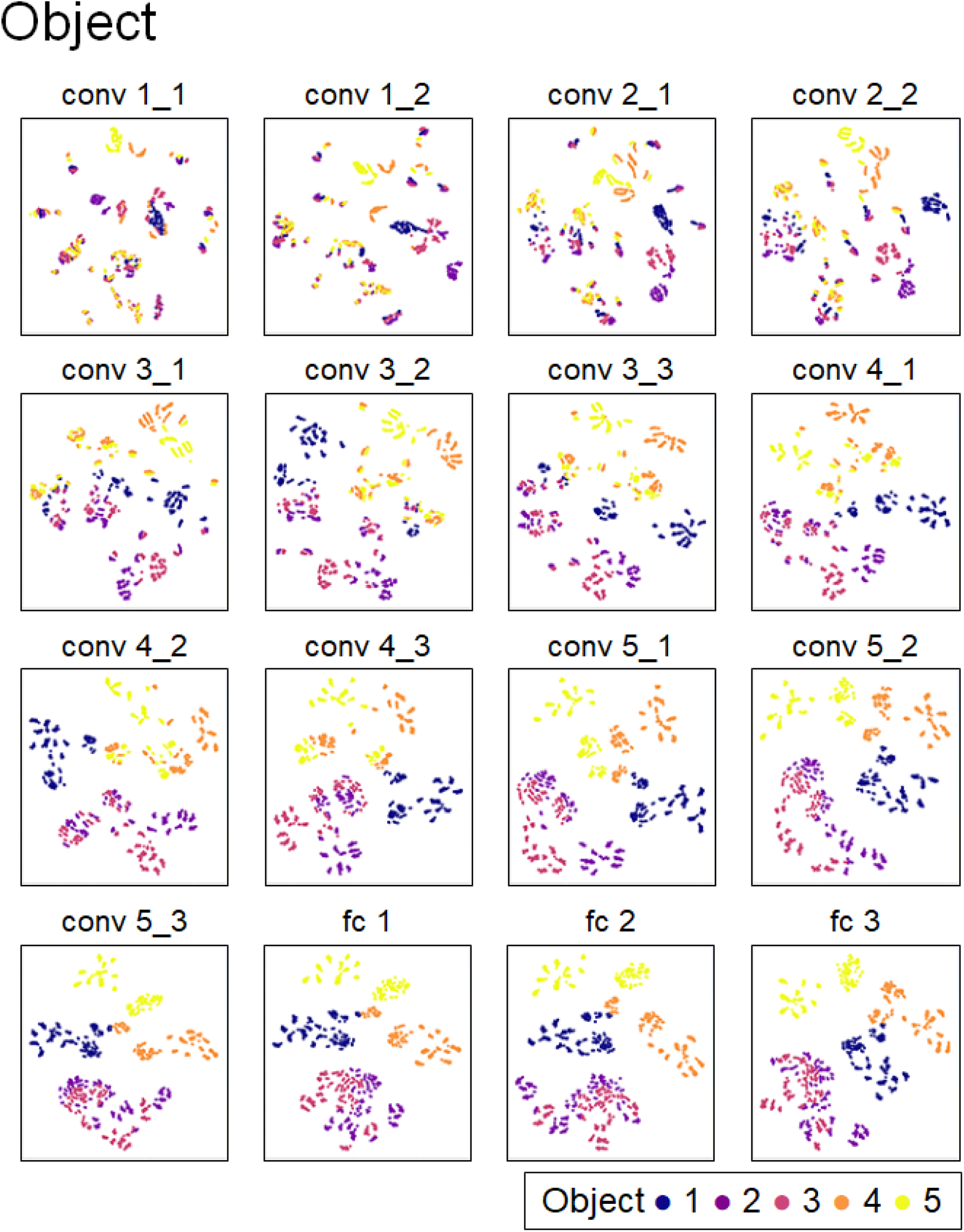
Embedding spaces of intermediate features of a deep neural network trained for object recognition. Results of all the 16 layers are shown with coloring different objects.

**Figure C4.**
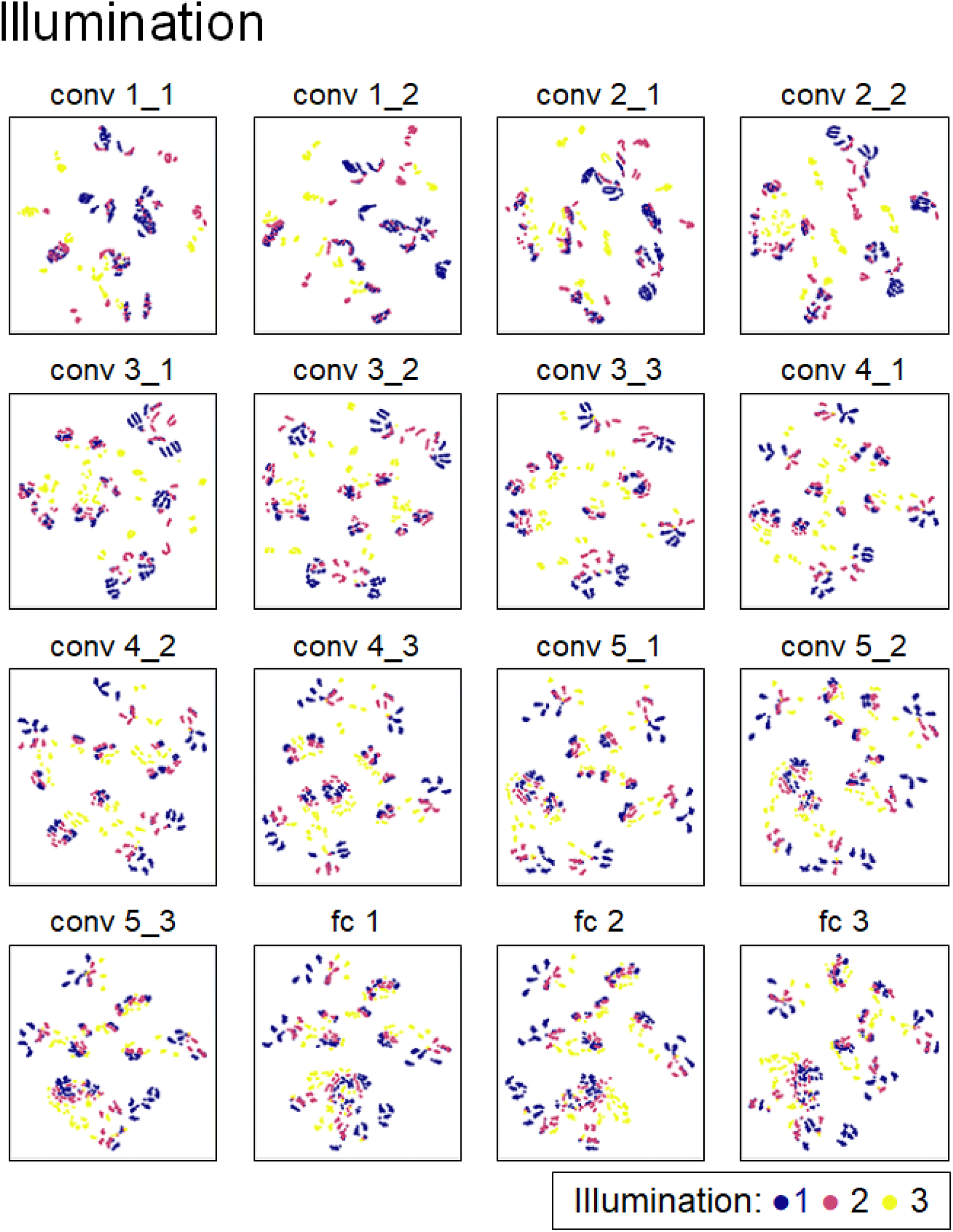
Embedding spaces of intermediate features of a deep neural network trained for object recognition. Results of all the 16 layers are shown with coloring different objects.

## Notes

### Competing Interest Statement

The authors have declared no competing interest.

### Summary of Updates

Accepted at Journal of Vision

